# Beta oscillations and waves in motor cortex can be accounted for by the interplay of spatially-structured connectivity and fluctuating inputs

**DOI:** 10.1101/2022.06.15.496263

**Authors:** Ling Kang, Jonas Ranft, Vincent Hakim

## Abstract

The beta rhythm (13-30 Hz) is a prominent brain rhythm. Recordings in primates during instructed-delay reaching tasks have shown that different types of traveling waves of oscillatory activity are associated with episodes of beta oscillations in motor cortex during movement preparation. We propose here a simple model of motor cortex based on local excitatory-inhibitory neuronal populations coupled by longer range excitation, where additionally inputs to the motor cortex from other neural structures are represented by stochastic inputs on the different model populations. We show that the model accurately reproduces the statistics of recording data when these external inputs are correlated on a short time scale (25 ms) and have two different components, one that targets the motor cortex locally and another one that targets it in a global and synchronized way. The model reproduces the distribution of beta burst durations, the proportion of the different observed wave types, and wave speeds, which we show not to be linked to axonal propagation speed. When the long-range connectivity is anisotropic, traveling waves are found to preferentially propagate along the axis where connectivity decays the fastest. Different from previously proposed mechanistic explanations, the model suggests that traveling waves in motor cortex are the reflection of the dephasing by external inputs, putatively of thalamic origin, of an oscillatory activity that would otherwise be spatially synchronized by recurrent connectivity.

## Introduction

Neural rhythms are one of the most obvious feature of neural dynamics [1]. They have been recorded for more than 90 years in human [2] and they are a daily tool for the diagnosis of neurological dysfunction. Classic studies have shown that neural rhythms depend on neural structures and the behavioral state of the animal [3]. Examples include the theta rhythm, the gamma rhythm in the visual cortex or the fast 160-200 Hz cerebellar rhythm. However, in spite of their ubiquity and common diagnostic use, neural rhythms remain a somewhat mysterious feature of the brain dynamics. It remains to better understand how they are created and what they are a reflection of [4].

The beta rhythm consists of oscillations in the 13-30 Hz range. The motor cortex was found to be one of its main locations in early recordings in human subjects [5, 6]. It was recorded in cats during motionless focused attention, when fixating a mouse behind a transparent screen [7]. Subsequent studies in monkeys trained to perform an instructed-delay-task [8, 9] showed that beta oscillations develop in the motor and premotor cortex during the movement preparatory period and wane during the movement itself, in agreement with earlier observations in human [5, 6]. It was also noted that beta oscillations did not have a constant amplitude in time but rather appeared as synchronized bursts of a few cycles of oscillations, often synchronized on electrodes 1-2 mm apart. More recently, it was observed using multi-electrode arrays that beta oscillations can come as planar [10] or more complex [11, 12] waves propagating horizontally on the motor cortex. Our aim in the present paper is to develop a mechanistic framework for these observations and to compare it to available recordings in monkeys performing an instructed delayed reach-to-grasp task [12, 13]. The model is based on recurrent interaction between local populations of excitatory (E) and inhibitory (I) population of neurons coupled by longer range excitation. The local E-I module is well-known to exhibit oscillatory dynamics when the recurrent interaction between the E population is sufficiently strong. Long-range excitation between these local modules has been shown in previous works to synchronize the oscillatory dynamics of local E-I modules [14, 15]. Comparison with recordings leads us to propose that the whole network is close to the oscillation threshold and that fluctuating inputs from other structures, such as the thalamus, power bursts of beta oscillations. We show that such a spatially extended network submitted to both local and global external inputs exhibits waves of different types that closely resemble those recorded in monkey motor cortex. Our analysis makes testable, specific predictions on the structure of external inputs. More generally, it highlights the dynamical interplay between fluctuating inputs and intrinsic dynamics shaped by spatially structured connectivity, which is worth of further experimental investigation.

### Data and modelling assumptions

We ground our modeling by comparing it to recordings obtained during a delayed reach-to-grasp task in two macaque monkeys [12] that have been made publicly available [13]. As in other recordings in similar tasks [8, 9, 10], beta oscillations are prominent during the 1 s waiting time, the movement preparatory period, and wane during the movement itself, as shown in Fig. S1a. We thus focus on the recordings during the preparatory period in the following.

In order to develop a model of beta oscillations, assumptions have to be made on the source and mechanism of their generation. We explicit below our main ones and their rationale, since they differ from those made in some previous works.

Debate exists regarding the origin of beta oscillations in cortex. On the one hand, the basal ganglia display prominent beta oscillations during different phases of movement [16]. Therefore, one view is that beta oscillations in the motor cortex are simply conveyed to the motor cortex from other structures such as the basal ganglia. On the other hand, sources of beta oscillations have been identified in the cortex [17, 18]. That different cortical regions have intrinsic oscillation frequencies matching those of their prominent rhythms is also supported by TMS perturbation studies. Specifically, the premotor area/supplementary motor area 6 has been found to resonate at ∼ 30 Hz after stimulation by a short TMS pulse [19]. This seems difficult to explain if the rhythm frequency originates from a distant structure. We thus adopt the intermediate view, previously advocated by Sherman et al. [18], that beta oscillations are generated by recurrent interactions in the motor cortex but are strongly modulated by inputs from other structures, notably thalamic ones.

Assuming that beta oscillations originate within M1, the question arises of the main neuronal populations and recurrent interactions underlying rhythm generation. Two natural candidates are recurrent interaction between interneurons, or oscillations involving recurrent interactions between excitatory and inhibitory populations. Here, one important observation is that during the preparatory period of movement when beta oscillations are prominent, neurons do not fire periodically. As shown in Fig. S1b, isolated units from the recordings of Brochier et al. [13] display large CVs. Beta oscillations thus appears to be a collective phenomenon arising from sparse synchronization [20] of different non oscillating units. In such a regime, oscillations can emerge from recurrent interaction between interneurons, when inhibition is sufficiently strong. However, their frequencies are mainly controlled by the kinetics of synaptic transmission and tend to be in the upper gamma range or higher [21, 22, 23]. Lower frequencies arise in networks of E-I units, in which each of the two populations inhibits itself via a slower disynaptic loop. This leads us to assume that beta oscillations arise from sparsely synchronized oscillations arising from recurrent interaction between excitatory and inhibitory populations.

Our aim is to account for recordings obtained from 4 mm×4 mm multi-electrode arrays. Therefore, the spatial structure of the connectivity needs to be taken into account. We assume that each electrode records the activity of a local neuronal population. For computational tractability, we describe this population activity in a classic way by its firing rate [24, 25], but we choose a particular firing rate formulation (see *Methods*) that was shown to agree well with direct simulations of spiking network models in previous works [15, 26, 27]. We assume that neurons under different electrodes are mainly connected by excitatory connections and that the connection probability decreases with distance [28, 29].

## Results

### E-I modules connected by long-range excitatory connections exhibit oscillatory activity at beta frequency

The previous considerations lead us to study the model schematically depicted in Fig. 1a which generalizes one previously analyzed [15]. It consists of multiple recurrently coupled modules between excitatory (E) and inhibitory (I) neuron populations. These local modules are further coupled by long range excitatory connections. The different modules are placed on a square grid corresponding to the different electrodes. The decrease of synaptic connection probability with distance is described by the function *C*(**x**), chosen in the numerical simulations to be Gaussian with range *l* ∼ 1 mm. We have taken into account the relatively slow propagation speed (∼ 30 cm/s) along unmyelinated horizontal axons by introducing a delay proportional to distance, *D*|**x** − **y**|, between the activity *r*_*E*_ in excitatory population at location **y** and the corresponding postsynaptic depolarization in the E and I population at location **x**. The model is further detailed in *Methods*. We begin by examining the dynamics of this network when external inputs are constant and the number of neurons in each module is very large. The effect of time-varying inputs and fluctuations due to the finite number of neurons in each module are then addressed in the following sections.

**Figure 1:**
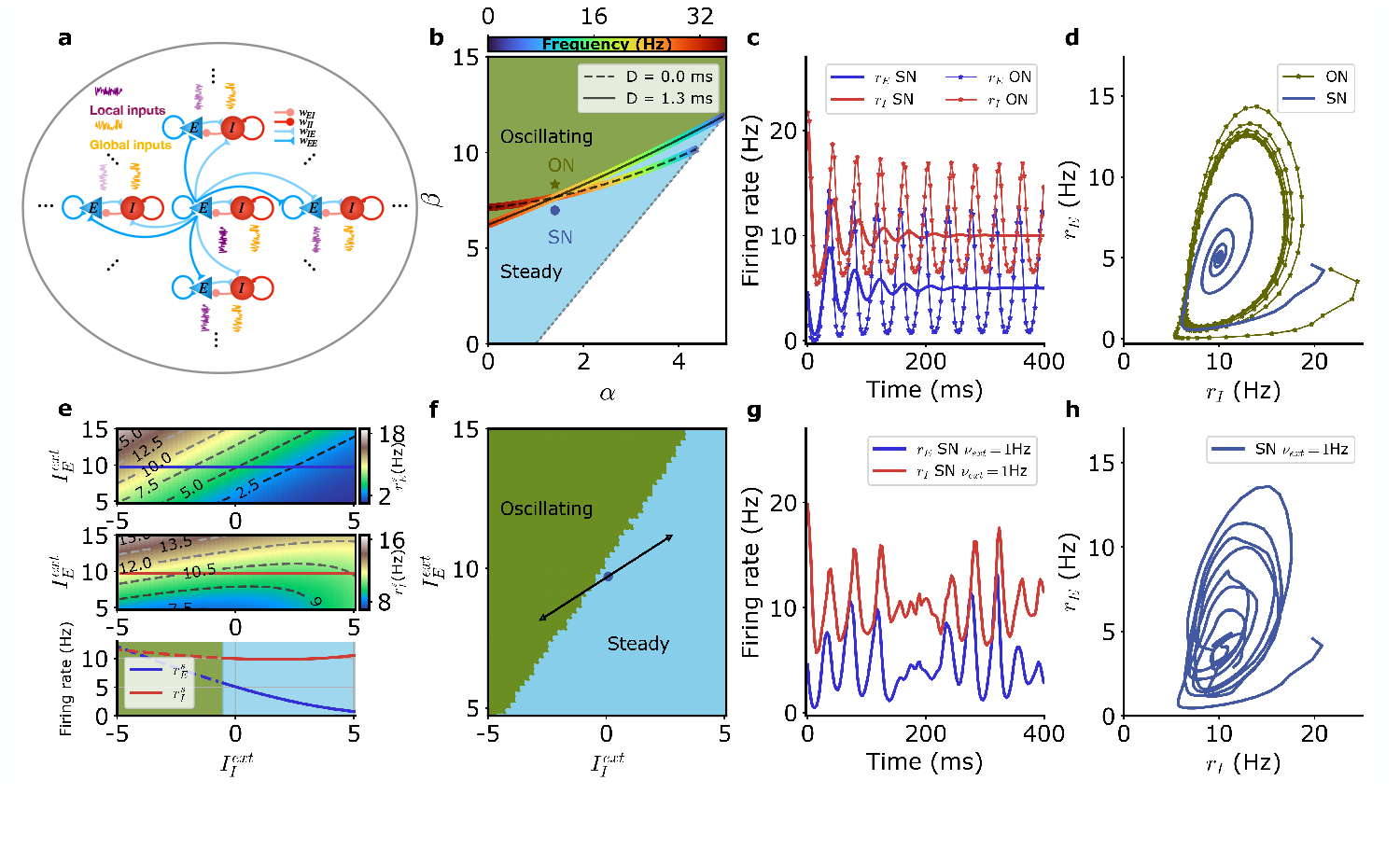
Model of neural network generating beta oscillations. (a) Schematic depiction of the model with excitatory neurons (blue), inhibitory neurons (red), independent external inputs on each module (“local”, purple) and inputs common to all modules (“global”, orange). (b) Different dynamical regimes for fixed firing rates of the excitatory and inhibitory populations 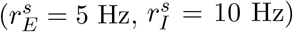, as a function of the strengths of recurrent excitation (*α*) and of feedback inhibition through the E-I loop (*β*) as defined in Eq. (32). The oscillatory instability line for *D* = 1.3 ms (solid black) and *D* = 0 ms (dashed black) and the line of ‘real instability’ (short-dashed black) are shown. Color around the oscillatory line indicates the frequency of oscillation at threshold at each point. (c) Time series of the firing rate for the E (blue) and I (red) module populations at the SN (thick lines) and at the ON (thin lines with symbols) points. E and I population activities become steady at the SN point and display regular oscillations at the ON point. (d) Same data as in (c) for SN (blue) and ON (green) parameters but with *r*_*E*_ plotted as a function of *r*_*I*_. (e) Firing rates of the E (top) and I (middle) populations for SN parameters when the external inputs are varied and (bottom) along the solid lines in top and middle panels when only the external input on the inhibitory population 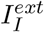 is varied (E blue, I red). The dashed parts of the line correspond to unstable steady states. (f) Different dynamical regimes as a function of the mean strength of external inputs for SN. The variation of the external inputs on each population is also shown (solid black line) when the strength of the external input is varied. (g) Example of E and I activity time traces when the external inputs vary in time along the solid dark line in (f). (h) Same data as in (f) but with *r*_*E*_ plotted as a function of *r*_*I*_. Model parameters correspond to SN and ON in Table 1.

The network of E-I modules, described by Eq. (8)-(17), has different dynamical regimes as a function of the synaptic weights *w*_*EE*_, *w*_*IE*_, *w*_*EI*_, *w*_*II*_ between the excitatory and inhibitory neural populations. This has been extensively studied in the rate model framework [30, 31] since the pioneering work of Wilson and Cowan [24]. In the particular case of the present model, the different regimes are depicted in Fig. 1b, generalizing previous results [15] to take into account the finite kinetics of synaptic current and propagation delays. Essential parameters controlling stability are the strength of recurrent excitation, the strength of feedback inhibition through the disynaptic E-I loop, and the strength of autoinhibition of interneurones on themselves, as respectively measured by parameters *α, β* and *γ* (Eq. (32)). With other parameter fixed, a steady firing rate state in which excitatory and inhibitory populations fire at 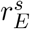 and 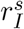 is unstable when the strength of recurrent disynaptic self-inhibition *β* becomes larger than a threshold value. Mathematical analysis (see *Methods)* provides the bifurcation line that separates the steady state regimes from the oscillatory ones. Its exact position depends on propagation delays as depicted in Fig. 1b for zero delay and for the propagation delay, here chosen, of 1.3 ms between neighboring modules, see also *Methods* Eq. (33). The frequency of oscillations that arise when crossing the bifurcation line at a particular point is also shown in Fig. 1b. The dynamics of the network is illustrated for two values of recurrent inhibition, one above (parameters ON, ‘oscillating network’) and one below (parameters SN, ‘steady network’) the bifurcation line point corresponding to beta oscillation frequency (∼ 20 Hz). As illustrated in Fig. 1c-d, for recurrent inhibition stronger than the critical value, the ON model oscillates regularly. For a lower value of recurrent inhibition (SN model), the firing rates of the excitatory and inhibitory populations steadily fire at constant rates. When perturbed away from these rates, the firing rates relax to their stationary values in an oscillatory fashion, transiently exhibiting beta oscillations.

**Table 1:**
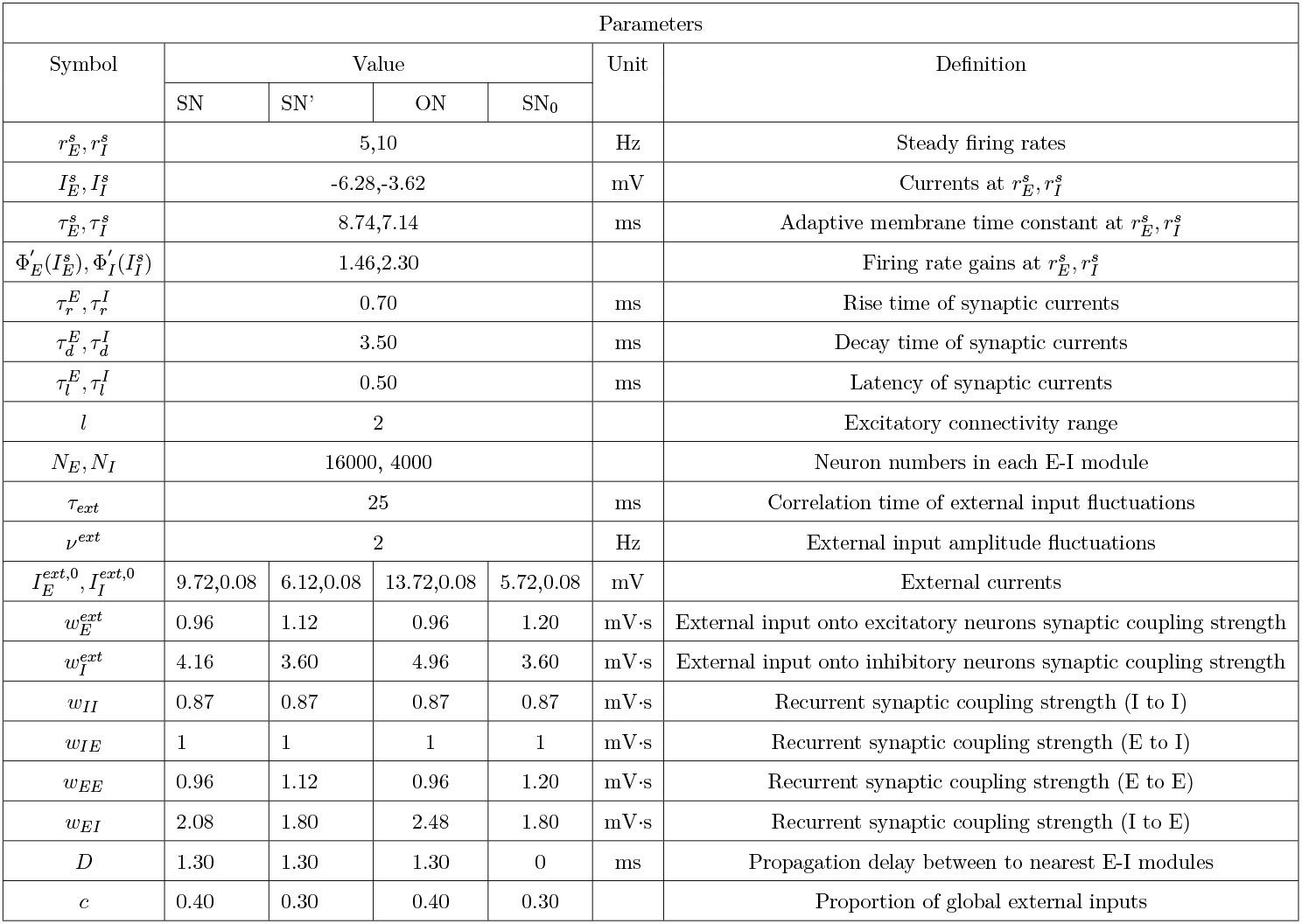
Parameter table.

For fixed synaptic coupling, the steady state firing rates of the excitatory and inhibitory neuronal populations depend on the strengths on the external inputs on those two populations. This is quantitatively illustrated in Fig. 1e, for synaptic parameters of model SN. Interestingly, when the external input on the inhibitory neurons is increased, their steady firing rate and the steady firing rate of the excitatory population both decrease for a large range of the external input current. (The firing rate of the inhibitory population starts to increase again for very large inputs, when the firing rate of the excitatory population tends to vanish.) This “paradoxical suppression of inhibition” [32] is typical of inhibition-stabilized networks and is observed in many cortical areas [33, 34] including motor cortex. It arises in the SN model because it is only when recurrent excitation is large, as measured by the parameter *α* that its frequency of oscillation is in the beta range (Fig. 1b). Namely all networks with *α >* 1 cannot stably fire at moderate rates without being stabilized by inhibition.

When the external input strengths are changed on the excitatory and inhibitory populations, the steady discharge rates of the two populations change accordingly. They can enter into the parameter regime where a steady discharge is actually unstable and is replaced by an oscillatory one, as shown in Fig. 1f. We consider in the present model excitatory inputs to the motor cortex coming from a distal area. The respective strengths of the external inputs, 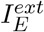 and 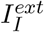, on the excitatory and inhibitory populations are proportional to the respective strengths of their synapses, 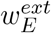 and 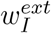. Thus, 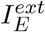 and 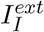 vary with the activity of the distal area on a line in the 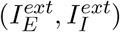 plane as depicted in Fig. 1f. The respective strengths of 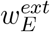 and 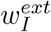 are taken in the model (Eq. (35)) such that the network enters the oscillatory regime when the external inputs are decreased as shown in Fig. 1f. This conforms to the early observation [5, 6] that external inputs suppress beta oscillations in motor cortex.

When the external input strength on the SN model varies in time, its operating point moves relative to the bifurcation line and correspondingly, beta oscillations are dampened at varying rates. This leads to waxing and waning of their amplitudes as shown in Fig. 1g-h. This is also true for the ON model which stands close to the bifurcation line and can move into the non-oscillatory regime when external inputs vary. These networks that operate close to the bifurcation line thus appear as promising candidates to describe the waxing-and-waning of beta oscillations seen in recording data. This leads us to try and compare beta oscillations produced in the model by inputs varying on an appropriate time scale, together with intrinsic fluctuations arising from the finite number of neurons in each module, to those seen in electrophysiological recordings.

### Fluctuation of external inputs, and finite-size stochasticity produces model LFP signals statistically similar to the recorded ones

Early reports [8] already stressed that beta oscillations were not of constant amplitude but modulated on different time scales. As confirmed in several later studies (see e.g. [35]), the average power of beta oscillations changes on a second time scale with different phases of movement e.g preparation, performance and post-performance periods. Within individual trials, the amplitude of beta oscillations fluctuates on much shorter time scales with brief bursts of elevated amplitudes of ∼ 100 ms duration. Examples of LFP spectrograms from two trials of [13] are shown in Fig. 2a-b and Fig. S2a-b), respectively for monkeys L and N. Whereas single trials exhibit short bursts of theta oscillations, trial-averaged spectrograms (Fig. 2c and Fig. S2c) and LFP power spectra (Fig. 2d and Fig. S2d) only show the average increase of oscillation in the beta band and its modulation on 1s behavioral time scale (Fig. 2c and Fig. S2c). A more quantitative characterization of the brief high oscillation amplitude events, the beta bursts, here taken to be the 75th percentile of the beta oscillation amplitude distribution (Fig. 2e and Fig. S2e, see also Fig. S3), is provided by their duration distributions shown in Fig. 2f and Fig. S2f.

**Figure 2:**
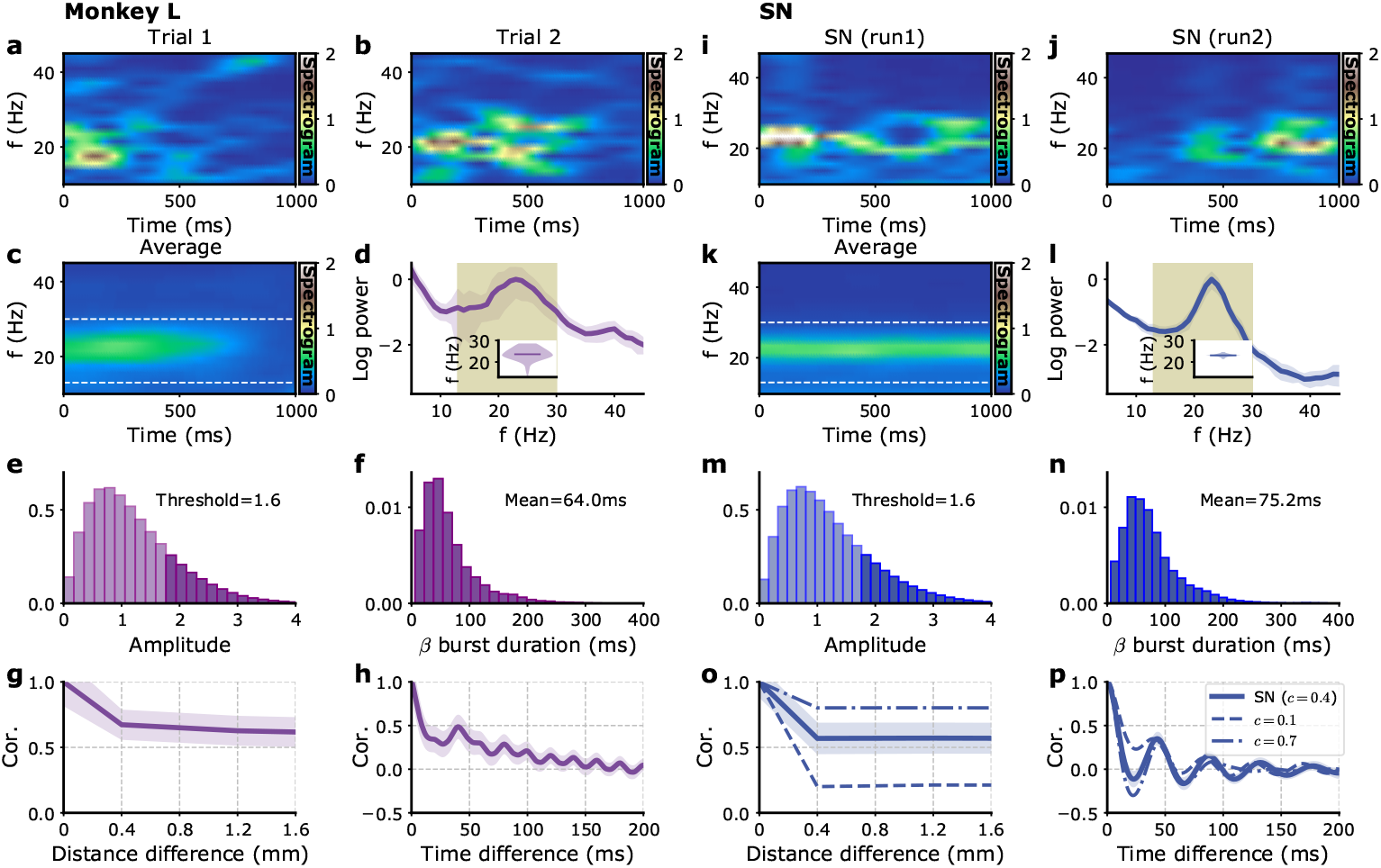
Single electrode recordings vs. model E-I module dynamics with fluctuating inputs. (a)-(h) Monkey L recordings [13], (i)-(p) Model simulations. (a)-(b) Two examples of single trials spectrograms of a single electrode LFP during the preparatory period (the CUE-OFF to GO period in Fig. S1). (c) Spectrogram averaged over different trials and different electrodes. (d) Power spectrum of single electrode LFP averaged over all electrodes. (e) Distribution of beta oscillation amplitudes. The amplitude of oscillation corresponding to the beta burst are shown in darker color. (f) Distribution of beta burst duration. (g) Average LFP cross-correlation between two electrodes as a function of their distance. (h) Auto correlation of single LFP time series averaged over trials and electrodes. (i)-(p) Corresponding model figures. (i)-(j) Two examples of single modules spectrograms of *I*_*E*_ 1 s time series. (k) Average *I*_*E*_ time series spectrogram. (i) Corresponding power spectrum. (o) Cross-correlations between different modules *I*_*E*_ times series as a function of their distance, for different *c* fractions of global (i.e shared) external inputs, *c* = 0.4 (solid), *c* = 0.1 (dashed) and *c* = 0.7 (dashed-dotted). (p) Auto correlation of single module *I*_*E*_ time series for the different fractions *c* in (o). Model parameters correspond to SN in Table 1.

Can the model introduced in the previous section account for these data? At least two sources can be considered for these transient bursts of oscillations fluctuating from trial to trial. The first one is that the number of neurons in each E-I module, the population of neurons in a 200 *μ*m neighborhood of each electrode, is finite, of about 2×10^4^ cells given the cell density in cortex. This by itself produces stochastic fluctuations in each module activity. These intrinsic fluctuations can be described in a simple way by adding to each neuronal population’s mean firing rate a stochastic component that is inversely proportional to the square root of its population size (see *Methods*) [15, 36]. The effects of these intrinsic fluctuations for models ON and SN (Fig. 1b) can be obtained both mathematically and through computer simulations. The results are displayed in Fig. S4 for varying numbers *N* of neurons per module around our estimated value *N* = 2 × 10^4^. For the parameters SN (Fig. S4a,b), the model LFP power spectrum amplitude grows with the number of neurons, but in the whole range *N* = 2 × 10^3^ − 2 × 10^5^, the power spectrum shape is invariant and accurately coincides with the one obtained mathematically by a linear analysis (Eq. (49)). Compared to the experimental LFP power spectra (Fig. 2d and Fig. S2d), the peak around beta frequencies is less pronounced; furthermore, the power spectrum misses the trough and enhancement for frequencies lower than beta frequency observed in the recordings. For the (oscillatory) ON model (Fig. S4j,k), the power spectra are more strongly peaked than the experimental ones, with in addition a notable peak at the harmonic frequency of ∼ 50 Hz coming from the non-sinusoidal shape of the oscillations (Fig. 1c,d). This indicates that in either the SN or the ON model, intrinsic fluctuations are not sufficient to account for the experimental spectra of Fig. 2d and Fig. S2d.

Besides intrinsic dynamical fluctuations, it is likely that fluctuations in the motor cortex arise from time-varying inputs coming from other areas. In absence of specific data, we model these as fluctuating inputs with a finite correlation time, more precisely as Ornstein-Uhlenbeck processes with a relaxation time *τ*_*ext*_ (*Methods*). The network dynamics also depend on the way these external inputs target the motor cortex. Are they essentially independently targeting local areas or do they target the motor cortex in a more globally synchronized manner? Again in absence of specific measurements, we suppose that the external inputs comprise a mixture of global inputs that provide a fraction *c* of the power, and independent local inputs that provide the remaining fraction 1 − *c* (see *Methods*).

With this simple description, external inputs are characterized by 3 parameters, their amplitude *ν*_*ext*_, their correlation time *τ*_*ext*_ and the fraction *c*. We determine them by comparison with the single electrode LFP trace and with the cross-correlation between the LFP traces of different electrodes.

The single electrode power spectra and the single electrode oscillation depend little on the fraction *c*, we thus consider them first (Fig. S4).

The inclusion of external fluctuations increases the lower frequency part of the power spectra. It also comparatively decreases the power at frequency higher than beta frequencies as soon as the fluctuation correlation time is in the few tens of millisecond range as shown for models SN (Fig. S4c-f) and ON (Fig. S4l-o). The amplitude of external fluctuations should be large enough for their effect to be of larger magnitude than the one produced by intrinsic fluctuations namely *ν*_*ext*_ *>* 0.2 Hz. For the SN model, it should also be small enough (*ν*_*ext*_ *<* 3 Hz), not to produce a secondary power enhancement at double the beta frequency as shown in Fig. S4c. In this range of amplitudes, the power spectrum shape is independent of the external fluctuation amplitude *ν*_*ext*_ after normalization, and corresponds to the mathematical expression (46) (Fig. S4d). For the ON model, a sharp secondary peak at twice the beta frequency is present at low external fluctuation amplitude (Fig. S4l). When the external fluctuations are stronger, this peak is blurred and becomes a wide power enhancement around twice the beta frequency for *ν*_*ext*_ ∼ 3 Hz (Fig. S4f,o), comparable to the one seen for the SN model at the same external fluctuation amplitude (Fig. S4c). The external amplitude fluctuations also produce the power spectrum trough and enhancement observed at lower frequencies in experimental recordings (Fig. 2d and Fig. S2d) when their correlation time is not too short (Fig. S4e-f,n-o).

Examination of the burst durations and amplitudes provides further information and constraints on the parameters. For both SN and ON models the burst mean duration grows with the input correlation time *τ*_*ext*_ (Fig. S4g,p) while it is weakly dependent on the input fluctuation amplitude *ν*_*ext*_ (Fig. S4i,r). For the SN model, the input fluctuation amplitude *ν*_*ext*_ does not strongly affect the burst amplitude and duration distributions (Fig. S4h-i). Their shapes are moreover close to the experimentally observed ones (Fig. (2e,f). On the contrary, for the ON model, the shape of the burst amplitude distribution strongly depends on the amplitude *ν*_*ext*_ (Fig. S4q). It is only for large *ν*_*ext*_ that the shape is close to the one obtained for the SN model and resembles the experimental one. The coincidence of the SN and ON distributions, as for the power spectra, is indeed expected when the fluctuations are large enough as compared to the difference between their reference parameters. Since the ON network appears potentially relevant only for large fluctuations, when the distinction before the two network set points is not meaningful, we focus on the following on a network at the set point SN, that is fluctuating around a non-oscillatory set point.

Fig. 2i-n and S2i-n, show the results of model simulations for the SN model with a relaxation time *τ*_*ext*_ = 25 ms and *ν*_*ext*_ = 2 Hz. Both LFP power spectra, and beta burst duration and amplitude distributions closely resemble the experimental ones.

Having reproduced the single electrode characteristics of the recording, we turn to the equal-time correlation between signals on different electrodes. As shown in Figs. 2g and S2g, the correlation between two neighboring electrodes is lower that the autocorrelation of a single electrode LFP, namely its variance, but is comparable to the correlation between more distant electrodes. Namely, part of the LFP signal is strongly synchronized between distant electrodes. This is not the case in the model if the external inputs on different modules are uncorrelated, i.e. for *c* = 0. At the other extreme, when *c* = 1, the signals are much more correlated than in the data. As shown in Fig. 2o, it is only when the external inputs are almost equally split between local and global that the model correlations are comparable to the experimental one. For monkey L (N), the value *c* = 0.4 (*c* = 0.3) is found to provide the best match. (Note that for the comparison with monkey N, we also use slightly changed synaptic couplings SN’ that lead to a slightly lower peak beta oscillation frequency in accordance with experimental data, see Table 1 for retained parameter values.) The autocorrelation of the single electrode LFP (Figs. 2h and S2h for monkeys L and N, respectively) is also well accounted for by the model (Figs. 2p and S2p for parameters SN and SN’, respectively).

Figs. 2 and S2 provide a comparison between the experimental data for the two monkeys and the model at points SN and SN’, respectively, with the determined characteristics of the external inputs. The model clearly accounts well for the data characteristics, as quantified by the power spectra, burst duration and amplitude distributions and correlation between different electrodes.

This leads us to investigate how the model compares to the data for other spatio-temporal characteristics of the LFP signals, namely traveling waves of activity of different types and their characteristics.

### Traveling waves of different types

Several groups have reported traveling waves on the motor cortex during the preparatory phase of the movement in instructed-delay reaching tasks. In the early study of Rubino et al. (2006) [10], planar waves of beta oscillations in multielectrode LFP recordings were described. Later works [11, 12] have classified the spatio-temporal beta oscillations recorded by multielectrode arrays into different states. Here, we use the data of [13] and closely follow the classification scheme proposed by these authors [12], distinguishing periods with planar or radial waves, as well as globally synchronized episodes and more random appearing states. The classification criteria into these four states are based on the instantaneous phase and phase gradient spatial distributions as precisely described in *Methods*. In short, negligible phase delays between electrodes characterize the synchronized state. In the other states, the oscillatory activities of some electrodes have significant phase differences and give rise to spatial phase gradients. The phase gradients are tightly aligned for planar waves, pointing inward or outward for radial waves, or are more disordered in the remaining “random” category. We apply the exact same analysis to the experimental recordings and our model simulation data (Fig.S5 and *Methods*).

Fig. 3a provides one example of a planar wave in monkey L recordings [13] with the narrow distribution of phase gradient directions shown in Fig. 3b. An example of a radial wave is shown in Fig. 3c with its outward pointing phase gradients shown in Fig. 3d. Analogous examples for monkey N are shown in Fig. S6.

**Figure 3:**
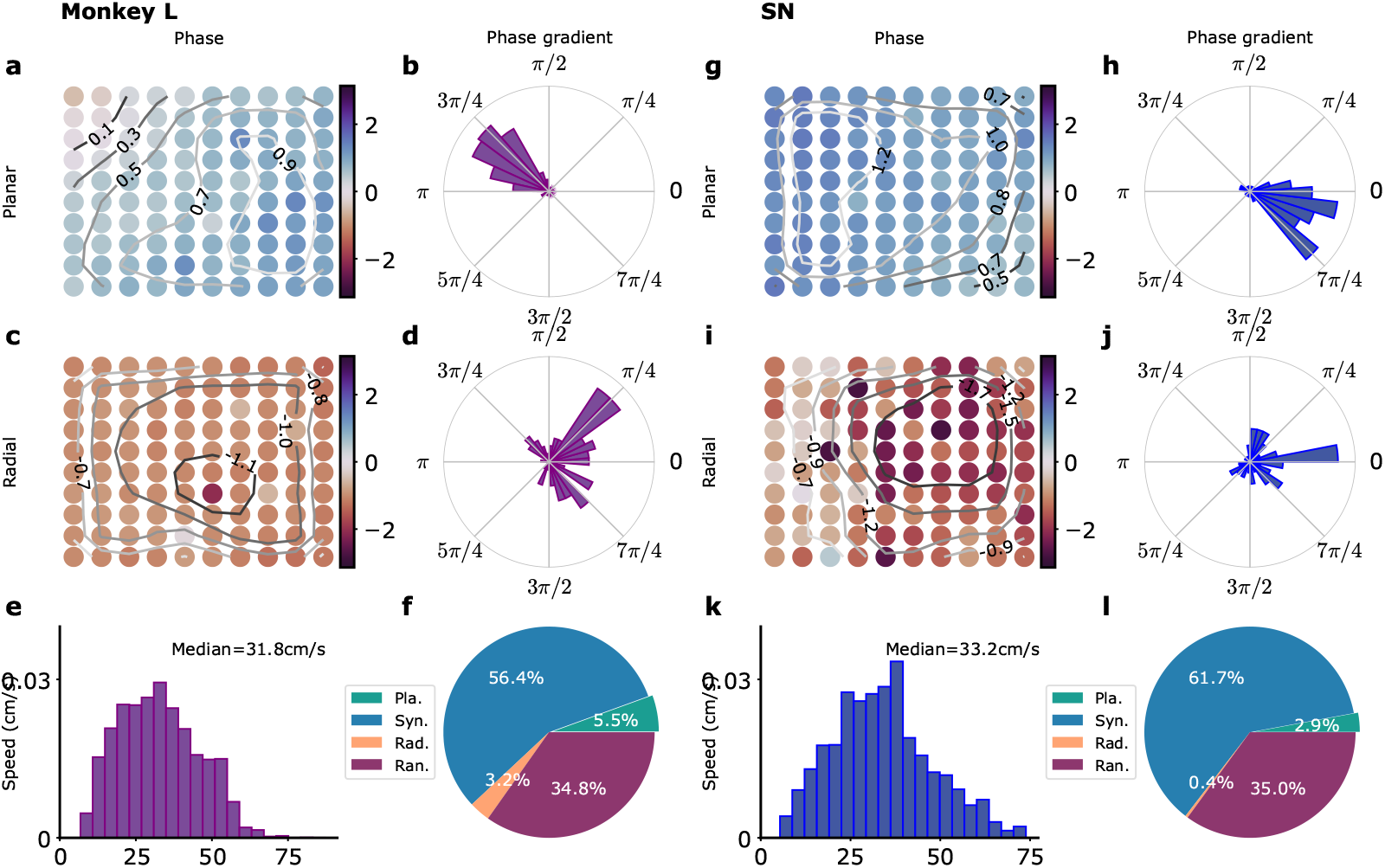
Waves in recordings and in model simulations. (a)-(f) Monkey L recordings [13], (g)-(l) Model simulations with SN parameters. (a) Snapshot of a planar wave. The phases of the LFP on the different electrodes are shown in color. Phase isolines are also shown (thin dark lines). (b) Distribution of phase gradients on the multielectrode array in (a) Note the high coherence of the phase gradients (*σ*_*g*_ = 0.61). (c) Example of a radial wave with LFP phases and isolines displayed as in (a). (d) Corresponding distribution of phase gradients (*σ*_*g*_ = 0.13). (e) Distribution over time and trials of measured planar wave speeds. (f) Proportion over time and trials of different wave types. (g)-(l) Corresponding model simulations. (g) Snapshot of a planar wave showing the *I*_*E*_ phases of different modules. (h) Distribution of *I*_*E*_ phase gradients (*σ*_*g*_ = 0.51) in (g). (i) Snapshot of a radial wave. (j) Distribution of phase gradients in (i) (*σ*_*g*_ = 0.13). (k) Distribution of planar wave speeds. (l) Proportions of different wave types.

For a traveling wave of oscillatory activity at frequency *f*, the local phase velocity is directly related to the local phase gradient ∇*ϕ*,

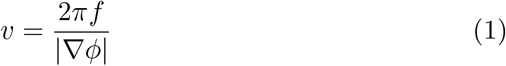

We refer to the average of *v* over electrodes simply as the wave speed following previous works [10, 11, 12]. The distribution of observed planar wave speeds is shown in Fig. 3e. The average planar wave speed is about 30 cm/s but it should be noted that the distribution includes also events with much smaller velocities. The proportion of the different wave types is depicted in Fig. 3f.

Can these data be accounted for by our model network? As discussed in the previous section, the single electrode LFP data and their correlations determine the correlation time (*τ*_*ext*_) of the fluctuating inputs and their repartition (*c*) between global and local inputs, but does not tightly constrain their amplitude. We thus performed a series of model simulations with different input amplitudes *ν*_*ext*_ for the SN model as summarized in Fig. S7a. The four wave states can be observed for most *ν*_*ext*_. Examples of planar and radial waves are provided in Fig. 3g-j. However, the proportion of the different wave types and the planar wave speeds greatly vary with *ν*_*ext*_ (Fig. S7). In the SN model, the input fluctuations promote oscillations. The stronger the amplitude *ν*_*ext*_, the more developed the oscillations in neural activity and the better they synchronize. Thus, increasing the noise amplitude first increases the proportion of synchronized activity and the propagation speed of planar waves. At higher noise amplitudes, the desynchronizing effect of noise counteracts its oscillation-promoting effect and the wave speed decreases with increasing input amplitude. This leads in the SN model for *ν*_*ext*_ in the range 1 − 2 Hz to a proportion of a planar wave states of a few percent with a planar wave speed of about 30 cm/s, as observed in the experimental recordings (Fig. 3e-f). As a consequence, the proportion of planar and radial waves as well as synchronized activity increases with *ν*_*ext*_. The effect of increasing the amplitude of the input fluctuations is quite different for the ON model (Fig. S7b). In this case, the oscillatory activity is self-generated and the oscillations between different modules are well-synchronized without fluctuating inputs. The local inputs tend to desynchronize the different modules and create oscillatory phase differences between them. The proportion of fully synchronized states thus decreases with the increasing *ν*_*ext*_. The speed of planar waves which is inversely proportional to the magnitude of phase gradients (Eq. (1)) also decreases. For an input fluctuation amplitude *ν*_*ext*_ = 2 Hz, the SN model agrees well with the distribution of observed wave speeds and the repartition between the different wave types for monkey L as shown in Fig. 3, although radial waves seem somewhat underrepresented in the model network. This is also the case for monkey N as shown in Fig. S6.

The mean speeds of planar waves in the recordings and in the model, are of a few tens of cm/s (Fig. 3e,k and Fig.S6e,k) which is comparable to the propagation speed along non-myelinated horizontal fibers. One might thus think that both are tightly linked. This is actually not the case. An equally good agreement with the recording data can be found in the model in the absence of propagation delay, i.e. for *D* = 0. The corresponding wave pattern statistics are shown in Fig. S8, and agree as well as the model with delay with the data for monkey L (Fig. 2 and 3).

### Synaptic connection anisotropy and planar waves propagation direction

Our model network provides a satisfactory description of LFP correlations and wave statistics when averaged over directions in the data of [13]. The long-range synaptic connection probability that we implemented decreases with distance and is independent of orientation. With this choice, we checked that the direction of observed planar waves does not significantly depend on orientation as shown in Fig. 4a-c (namely the anisotropy created by our choice of a rectangular grid is weak). However, it was reported by [10] that waves tend to propagate along preferred axes on the cortical surface with respect to sulcal landmarks and that the orientation of the preferred axes of propagation also vary between different regions, e.g. between primary motor cortex and dorsal premotor cortex. For a given preferred axis, propagation in both directions was observed, e.g. both rightward and leftward propagating waves were recorded. In order to investigate the possible origin of this observation, we assessed the effect of anisotropic connectivity in our model. We thus simulated a modified version of the model in which the connectivity was of twice longer range in the y-direction than in the x-direction, as illustrated in Fig. 4d. This did not change much the distribution of the observed wave types (Fig. 4e). In contrast, Fig. 4f shows the measured distribution of the directions along which planar waves propagate when connections are asymmetric. The fractions of waves propagating along the x and y axis are clearly different. There is a large predominance of waves observed to propagate along the x-axis, namely along the axis where the connectivity is shorter range. Longer-range connectivity promotes longer-range synchronization along the y-axis and synchronized states. The weaker connectivity along the x-axis allows for the formation of the larger phase gradients along the x-axis and planar waves.

**Figure 4:**
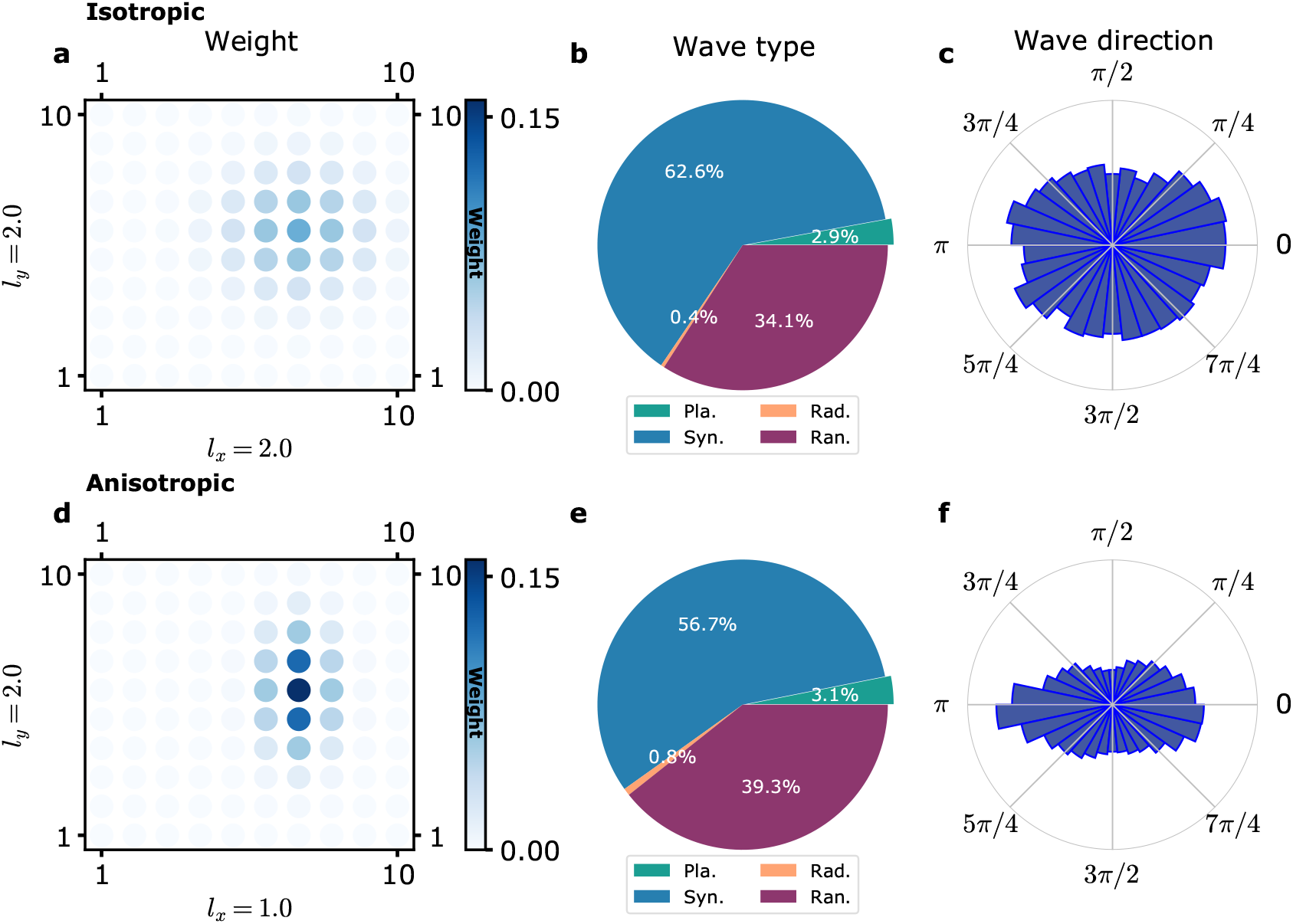
Influence of connectivity anisotropy on planar wave propagation direction. (a)-(c) Model simulations with an isotropic connectivity as in the previous figures. (a) The function *C*(**x, y**) is shown (with **x** arbitrarily chosen at position (7,6)). (b) Proportion of different wave types. (c) Distribution of propagation directions of planar waves. (d) The chosen anisotropic function *C*(**x, y**) is shown (with **x** again chosen as (7,6)). (e) Proportion of different wave types. (f) Distribution of propagation directions of planar waves. The planar waves predominantly propagates along the x-axis, the axis along which *C* decreases the fastest.

## Discussion

We have proposed and studied here a simplified model of the motor cortex based on local recurrent connections coupling excitatory and inhibitory neuronal populations together with long-range excitatory connections targeting both populations. The well-known oscillatory instability of coupled E-I networks is not qualitatively modified by the presence of long range connections and the model displays an instability leading to sustained oscillatory behavior. These oscillations take place at beta frequency for adequate synaptic parameters. Comparison with available data has led us to conclude the motor cortex operates close to this beta oscillatory instability line under strong fluctuating inputs from other areas. This constantly displaces the operating point of the dynamics and lead to waxing and waning of beta amplitude oscillations. Analysis of this behavior in time and across electrodes has led us to infer the characteristics of the appropriate external inputs. The observed power enhancement on the low frequency-side of the beta peak requires that their time correlation is not too short on a 10 ms scale while the duration of the observed oscillation bursts requires than it is not too long on the same time scale. In an analogous way the external input amplitude should be, on the one hand, sufficiently large to significantly modify the LFP characteristics above the background of intrinsic local fluctuations arising from the finite number of neurons sampled by each electrode. On the other hand, it should be small enough to preserve the harmonicity of beta oscillations, as indicated by the absence of a significant secondary 50-60 Hz peak in the LFP spectra. The spatial correlation of beta oscillations at the millimeter distance scale, has furthermore led us to suggest that the motor cortex receive inputs that target local areas but also synchronous inputs that target it more globally. Under these conditions, traveling waves of different kinds are observed that resemble those recorded in vivo both in their repartition between the different wave types and their speeds of propagation. Characterization of the inputs onto motor cortex during movement preparation is needed to test these theoretical predictions.

Propagating waves of neural activity have been observed in different contexts [37, 38]. Some are unrelated to neural oscillations such as sub-threshold waves [39] or propagation of spiking activity in visual cortex [40]. Others are based on oscillatory activity with proposed mechanisms relying on a gradient of frequencies [41] or structural sequences of activation [42]. The traveling waves and the mechanism here proposed to underlie them are quite different. They are obviously based on the existence of oscillations, since the traveling waves are a reflection of different phases of oscillation at different positions on motor cortex. The model we have developed here includes structural connectivity which tend to synchronize oscillations at different positions, but the stochastic external inputs play an essential role in creating dephasings between different locations. The wave propagation speeds are found to be rather low and distributed in the few tens of cm/s. The mean speed of about 30 cm/s is comparable to the propagation speed along non-myelinated horizontal fibers. However, we have found that it is actually independent of it, a comparable distribution of traveling speeds is found in a model with no propagation delays. This stands, for instance, in sharp contrast with a recent model of the propagation of spiking activity in visual cortex [43] which does not involve oscillatory activity but also relies on external inputs. In the present context, the observed speeds of the oscillatory waves of activity are set by the oscillatory frequency, and by the dephasings produced by the external inputs in the recorded area.

We have proposed that external inputs have both local and global components. The existence of a global component appears consistent with the presence of global inhibition in motor cortex during movement preparation [44]. The model “external inputs” could represent direct inputs from different neural structures, including frontal and parietal cortex and thalamus [45]. The known thalamo-cortical connectivity [46] makes the thalamus a privileged candidate for the origin of, at least, part of these external inputs. Indeed the described diffused connectivity from calbindin-positive matrix neurons could be the source of our global inputs while core parvalbumin-positive neurons could be the source of our local inputs. This requires further experiments assessing the origin of synaptic inputs and their influence on waves and wave types. As far as recurrent connectivity is implied, we have suggested that the observed anisotropy of wave propagation is linked to the anisotropy of long-range connections, with the specific prediction that traveling waves of oscillatory activity are predominantly observed orthogonal to the longest connection axis. This also needs to be tested in further experiments.

Finally, the role of traveling waves during movement preparation remains to be clarified. We have shown that a stochastic description of the inputs to the motor cortex is sufficient to account for the recordings. However, the produced traveling waves are correlated with the particular inputs that the motor cortex receives. Indeed, traveling waves have been shown to carry information about the subsequent movement [10]. Movement preparation is presently viewed, in a dynamical system perspective, as taking place in a subspace orthogonal to the dynamics of the movement itself and producing the proper initial condition for it [47, 48, 49, 50, 51]. Beta oscillations and their associated timescale are however absent from this description. Further work is needed to refine it and include beta oscillations and waves and obtain a more comprehensive description of motor cortex dynamics.

## Methods

### Recording data

We analyzed the data that was made publicly available and described in detail in [13]. We content ourselves here to describe their main features, for the convenience of the reader. The data were obtained from two macaque monkeys (L and N) trained to perform one of 4 movements at a GO signal. In brief, in each trial, the animal was presented for 300 ms with a visual cue that provided partial information on the movement to be performed. It had to wait for 1 s, before the GO signal that also brought the missing information about the rewarded movement. The different phases of a trial are depicted in Fig. S1.The neural activity was recorded during the task with a 10 × 10 square multielectrode Utah array, with neighboring electrodes separated by 400 *μm*, implanted in the primary motor cortex (M1) or premotor cortex contralateral to the active arm. We make use of two published recording sessions (one for monkey L of 11:49 min/135 correct trials, one for monkey N of 16:43min/141 correct trials ) as fully detailed in [13].

### Data analysis

An overview of our data analysis protocol is provided in Fig. S5. The different steps are detailed below.

### Data filtering

The index of each electrode in the data (Fig. S3) was matched to the (*x, y*)-position of the electrode in the data which we use in the following. The few missing electrode signals were replaced by the average of the neighboring electrode signals. The followings steps were applied to the so obtained *N* = 100 electrode signals. The LFP signal of each electrode was band-pass filtered in the 13–30 Hz range using a third-order Butterworth filter (`signal.filtfilt` function of the Python package `scipy`). The filtered signal of each electrode was then z-score normalized and Hilbert-transformed, using the `scipy` Hilbert transform function `signal.hilbert` to obtain its instantaneous amplitude *A*_*xy*_(*t*) and phase Φ_*xy*_(*t*) (Fig. S3). The instantaneous maps *A*_*xy*_(*t*) and Φ_*xy*_(*t*) were used for beta bursts and wave pattern detection, respectively.

### Beta bursts analysis

For each amplitude signal *A*_*xy*_(*t*), the percentiles of the amplitude distribution were determined [52]. The beta burst threshold was set at the 75*th* percentile. A burst onset was defined as a time point at which the analytical amplitude exceeded the threshold and its termination as the earliest following time point at which the amplitude fell below the threshold. The time difference between these two time points was defined as the burst duration. The amplitude of a burst was defined as the average amplitude during the burst duration. The procedure is illustrated in Fig. S3.

### Spatio-temporal pattern analysis and wave classification

The spatio-temporal patterns of oscillatory activity were classified with the help of the phase map Φ_*xy*_(*t*) following the steps of [12] with minor modifications.

First, the phase gradient map Γ_*xy*_(*t*) and its normalized version, the phase directionality map Δ_*xy*_(*t*), were obtained. Specifically the phase gradient map was obtained by averaging the phase differences between a point and its existing next and next-nearest neighbors along the x and y axis,

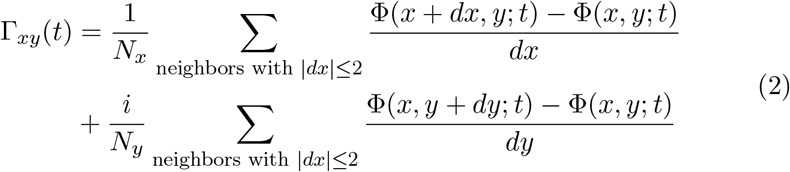

where the vector has been written as a complex number and *N*_*x*_, *N*_*y*_ count the actual number of neighbors along x and y respectively (*N*_*x*_ + *N*_*y*_ may differ from 8 for electrodes near the array boundaries). Normalization of the gradients to unit length serves to produce the “phase directionality map” [12],

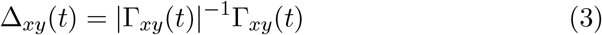

The alignment of the unit vectors Δ_*xy*_(*t*) at time *t* was quantified using the norm *σ*_*g*_ of their mean, the “circular variance of phase directionality” [12], with

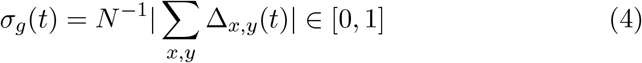

with *N* the total number of electrodes. Well-aligned Δ_*xy*_(*t*) were classified as planar wave patterns using the criterion *σ*_*g*_(*t*) *>* 0.5, and provided the first category of the classification.

The remaining patterns were further classified using additional measures. Patterns were next categorized as “radial waves” (or not) by considering critical points (i.e. maxima, minima or saddle points) of the phase map Φ_*xy*_(*t*) [11]. A smoothed phase gradient map, the “gradient coherence map”, was defined following [12],

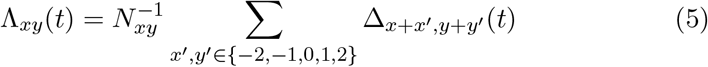

(Note that the sum is taken over existing neighboring electrodes, the number of which is given by *N*_*xy*_ ≤ 9.) Minima, maxima, and saddle points were obtained by locating the sign changes in the local gradient coherence map [53]. Maps with exactly one critical point were classified as radial waves, thus providing the second category of our classification (these patterns could be subdivided in further subcategories [11] but this was not pursued).

The synchronous patterns were finally identified using the “circular variance of phases” [12], *σ*_*p*_, analogous to the well-known Kuramoto order parameter,

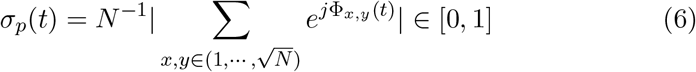

with *N* the total number of electrodes in the array. Patterns with *σ*_*p*_(*t*) *>* 0.85, indicating tight synchronization of the oscillations of all electrodes, were classified as “synchronized” the classification third category. All remaining patterns were assigned to the “random” fourth category.

### Model

The motor cortex is described as a collection of recurrently connected excitatory (E) and inhibitory (I) neuronal populations of linear size ∼ 400*μm*, comparable to the one of a cortical column. These E-I modules are connected by long-range excitatory connections with a connection probability decaying with the distance between modules. The neural activity of each E-I module is represented at the level of its excitatory and inhibitory neuronal populations in the rate-model framework [24, 25]. This eases numerical simulations that would otherwise be extremely demanding for a comparable spatially structured network of a few hundred modules with a few tens of thousand spiking neurons each. Additionally, the rate model description also eases analytical computations. We choose a rate model description with an adaptive time scale [26, 27]. It was shown to quantitatively describe networks of stochastically spiking Exponential Integrate-and-Fire (EIF) neurons, either uncoupled and receiving identical noisy inputs [26], or coupled by recurrent excitation [27], as well as sparsely synchronized oscillations of recurrently coupled spiking E-I module of EIF neurons [15]. The adaptive timescale rate model describes the activity of a population of EIF neurons as:

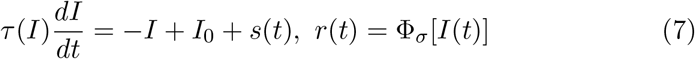

with Φ_*σ*_[*I*] the f-I curve of an EIF neuron with white noise current input of mean *I* and strength *σ* (see [15] for a detailed description). The time scale *τ* (*I*) is referred to as ’adaptive’ because it varies with the current *I*. The function *τ* (*I*) is chosen here, as in [15] and precisely described there, to best fit the response to oscillatory inputs in a wide frequency range of 1Hz-1kHz, of an EIF neuron subjected also to a white noise current of mean *I* and strength *σ*. The tabulated functions Φ_*σ*_[*I*] and *τ* (*I*) are given in [15] and are also provided here as *Source data* for the convenience of the reader.

We generalize the rate model description (7) to a two-dimensional network of E-I modules coupled by long-range excitation,

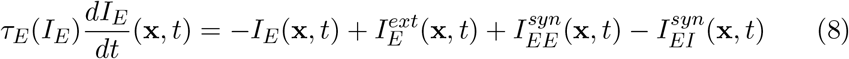

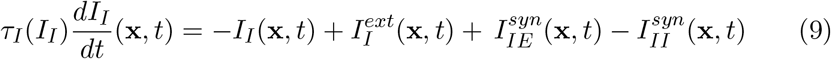

where the index **x** denotes the module position on a rectangular grid. The firing rates *r*_*E*_(**x**, *t*), *r*_*I*_(**x**, *t*) are related to the currents *I*_*E*_(**x**, *t*), *I*_*I*_(**x**, *t*) by,

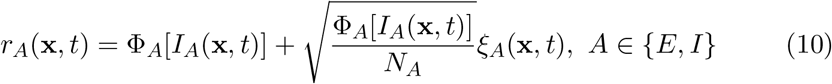

The *ξ*_*A*_(**x**, *t*) are independent unit amplitude white noises for each neuronal population in each module ⟨*ξ*_*A*_(**x**, *t*)*ξ*_*B*_(**x**^**′**^, *t*^′^)⟩ = *δ*_*A,B*_*δ*_**x**,**x′**_ *δ*(*t* − *t*′) and Ito’s prescription is used to define Eq. (10) [15]. These stochastic terms account, in the firing rate framework, for the fluctuations due to the finite numbers *N*_*E*_ of excitatory neurons and *N*_*I*_ of inhibitory in each E-I module [36]. For the numerical computations, both Φ_*E*_ and Φ_*I*_ are taken equal to the function Φ_*σ*_[*I*] provided in the *Supplemental Material*.

The currents 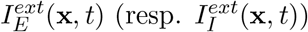 represent inputs external to the motor cortex, possibly position and time-dependent, targeting the excitatory (resp. inhibitory) population of the E-I module at position **x**,

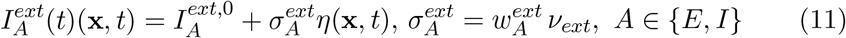

We model the fluctuations of external inputs as stochastic O-U processes with global and independent components,

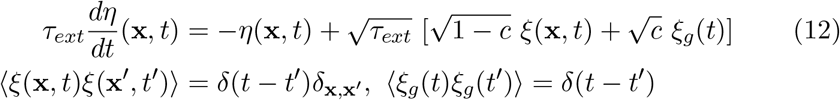

The currents 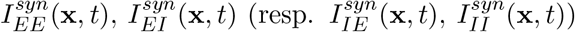 represent the recurrent excitatory and inhibitory inputs on the excitatory (resp. inhibitory) population of the module at position **x**. With our normalization choice, *ν*_*ext*_ has the dimension of a frequency and can be interpreted as the discharge rate ampltude of the external inputs. The recurrent inputs depend on the firing rates of the neuronal populations of the different motor cortex modules and on the kinetics of the different synapses. Namely,

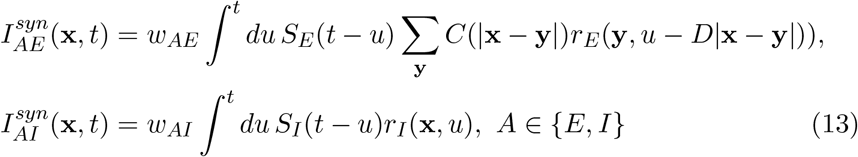

The kinetic kernels *S*_*E*_(*t*), *S*_*I*_(*t*) include synaptic current rise times, 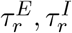, decay times, 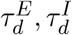, and latencies, 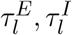,

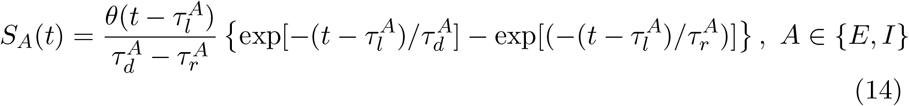

where *θ*(*t*) denotes the Heaviside function, *θ*(*t*) = 1, *t >* 0 and 0 otherwise. The kernels *S*_*A*_, *A* ∈ {*E, I*} are normalized such that:

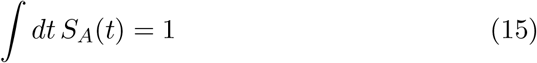

We have supposed, for simplicity, that the kinetics of the synaptic currents depend on their excitatory or inhibitory character but not on the nature of their post-synaptic targets (e.g. 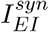 and 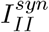 have the same kinetics). Instead of using the kinetic kernels *S*_*A*_(*t*), *A* ∈ {*E, I*}, one can equivalently compute the synaptic currents by introducing supplementary variables *J*_*AE*_, *J*_*AI*_,

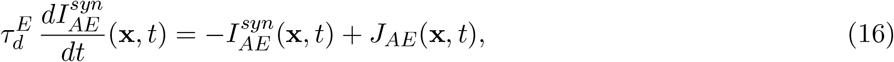

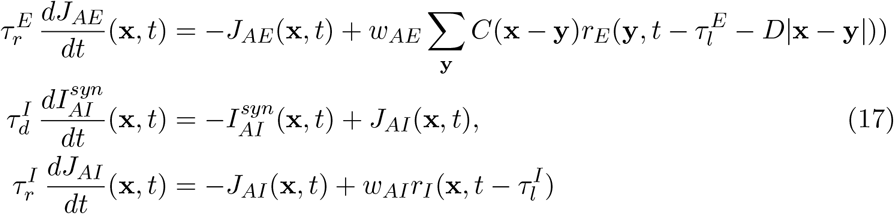

In Eq. (13) or (16), the probability that an excitatory neuron at position **y** targets a neuron at position **x** is represented by the function *C*(**x** − **y**) normalized such that,

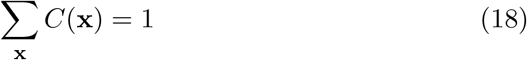

The delay, *D*|**x** − **y**| accounts for the signal propagation time along horizontal non-myelinated fibers in the motor cortex. The mathematical expressions below are written for a general function *C*(**x**). For all the numerical computations, except those shown in Fig. 4, the following isotropic Gaussian function is taken,

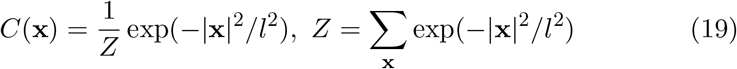

In Fig. 4, we consider instead the anisotropic function,

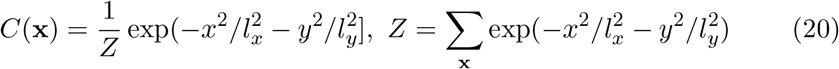

with **x** = (*x, y*).

### Theoretical analysis

Our analysis starts by considering the deterministic version of the model described by Eq. (8)-(17). That is, the limit *N*_*E*_ → ∞, *N*_*I*_ → ∞, in which the amplitudes of the stochastic terms vanish in Eq. (10). All module populations are supposed to fire steadily in time at constant rates 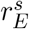 and 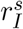. This requires external inputs that are constant in time, i.e. with 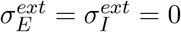 (Eq. (11)), and of specific magnitudes that we first determine.

We then assess the stability of this steady state. This provides the bifurcation diagrams of Fig. 1. When the steady state is stable, we compute the effect of fluctuations arising both from the finite number of neurons in each E-I module and from the time-variation of the external inputs. This provides the analytic curves for the power spectra and correlations of currents, shown in Fig. 2, S2, and S4.

### Steady state

We consider first the steady state of the deterministic network in which excitatory populations and inhibitory populations are firing at constant rates 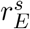 and 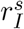, independently of time and module position, with

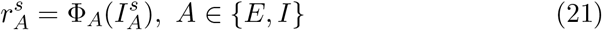

In this state, the synaptic currents in the different populations are obviously also independent of time and position. Given the normalization conditions (Eq. (15), (18)) they simply read,

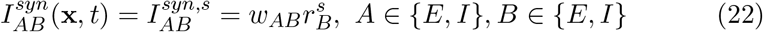

From the model definition (Eq. (8), (9)), these firing rates are produced by constant external inputs of magnitudes,

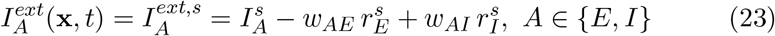

### Bifurcation lines

The stability of the steady firing state can be assessed by imposing the constant external currents (Eq. (23)) and computing the dynamics of small perturbations around the steady state. To this end, we linearize Eq. (8)-(17) and look for solutions that are oscillatory in space and exponential in time,

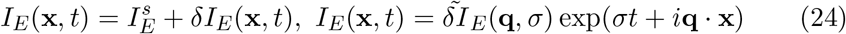

with similar expansions for the other variables. Substitution in the explicit formulas (Eq. (14)), or in the corresponding differential equations (Eq. (16,17)), provides the expression of the synaptic current in term of the module activities,

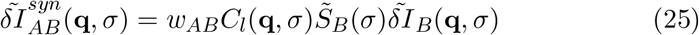

where 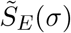 and 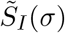 are the Laplace transforms of *S*_*E*_(*t*) and *S*_*I*_(*t*),

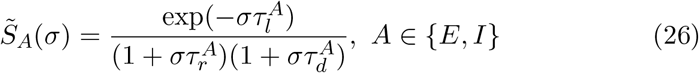

Substitution of Eq. (25), in the linearized Eq. (8, 9) gives

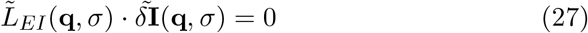

The matrix 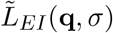 reads,

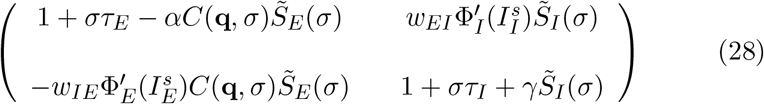

where the function *C*(**q**, *σ*) is the Fourier transform of the coupling function with propagation delays taken into account,

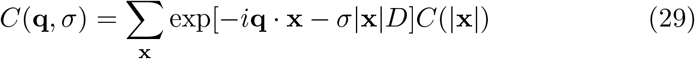

In Eq. (28) and in the following, the short-hand notation *τ*_*E*_ (resp. *τ*_*I*_) is used for 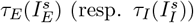. The existence of a non-trivial solution of Eq. (27) requires that the determinant, *W* (**q**, *σ*), of the matrix 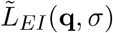 vanishes, i.e.

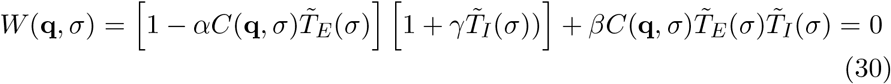

with the functions 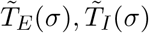, defined by,

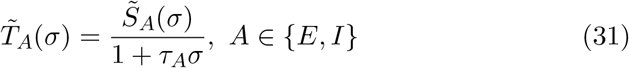

The constants *α* and *γ* respectively measure the gain of monosynaptic recurrent excitation and inhibition while *β* measures the gain of disynaptic recurrent inhibition,

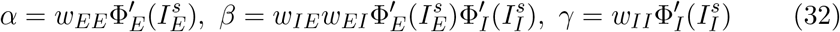

where 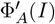 denotes the derivative of Φ_*A*_ with respect to *I*.

The oscillatory instability, or “Hopf bifurcation”, line, corresponds to the parameters for which the growth rate is purely imaginary, *σ* = *iω*. It is obtained in parametric form, with *α* and *β* as functions of the frequency *ω* (and of the recurrent inhibition *γ*) by separating the real and imaginary parts of Eq. (30) and solving the resulting linear equations, for *α* and *β*. This gives,

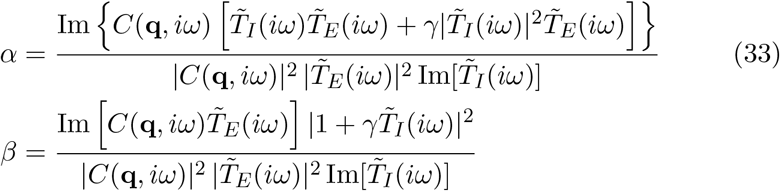

The instability first appears at long wavelengths, namely at **q** = 0 on a large enough lattice. The expressions (33) with **q** = 0 have been used to draw the diagram of Fig. 1b.

Besides this oscillatory instability, there is a possible loss of stability towards a high firing rate state. It is obtained when the growth rate *σ* changes with parameter variation from being real negative to real positive i.e. for parameters such that *W* (**q**, 0) = 0. Since at the ’real’ instability threshold, the instability growth rate vanishes, the instability line is independent of the synaptic current kinetics. One indeed checks from Eq. (26, 31) that 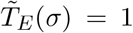. Thus, Eq. (30) gives for the real instability line for a given wavevector **q**.

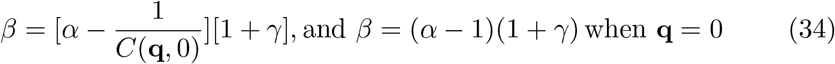

Since *C*(**q**, 0) *<* 1 when **q** does not vanish, the instability appears first at **q** = 0 with *C*(**0**, 0) = 1 when all the modules are in the exact same state.

The expressions (33, 34) with **q** = 0 have been used to draw the diagram of Fig. 1b. The diagram of Fig. 1f is similarly obtained by computing the external inputs corresponding to different steady discharges, for fixed synaptic parameters, and assessing the stability of these states from their position with respect to the stability lines (33) and (34). We have taken the synaptic strength of external inputs such that the network operating point crosses the oscillatory bifurcation line when the external input amplitude varies,

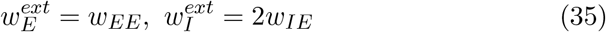

### Auto and cross-correlations of module activities and power spectrum

We consider the network described by Eq. (8, 9) with the kinetics of the synaptic currents given by Eq. (14). We include the stochastic effects arising from the finite number of neurons in each module by using the stochastic description (Eq. (10)) of the instantaneous firing rates of the excitatory and inhibitory module neuronal populations. We also include external input fluctuations as described by Eq. (11,12).

We treat these two kinds of stochastic effects as perturbations of the steady dynamics and fully characterize the stochastic dynamics of the network at the linear level.

Linearizing Eq. (8, 9) around their steady values gives,

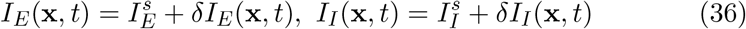

Translation invariance leads us to search for *δI*_*E*_ and *δI*_*I*_ in Fourier space,

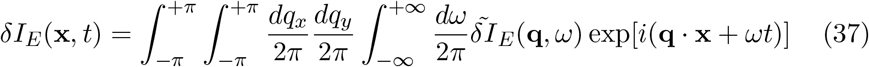

with an analogous expansion for *δI*_*I*_.

The linearized Eq. (8, 9) then read, with a vectorial notation

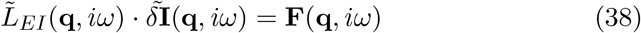

where the 2×2 matrix 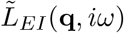 is given in Eq. (28). The two components of the stochastic forcing term **F**(**q**, *iω*) read,

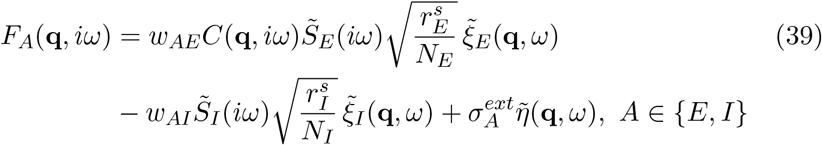

where 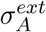 is given by Eq. (11) and (35). Solution of the linear system (38) provides the expression of the Fourier components of the currents. For the excitatory current, one obtains,

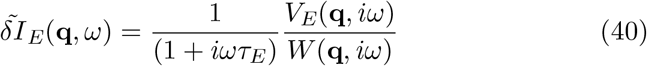

with *W* (**q**, *iω*) defined in Eq. (30) and *V*_*E*_(**q**, *iω*) given by,

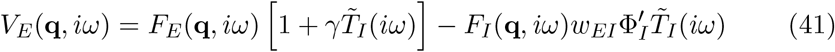

The Fourier components of the input fluctuations read,

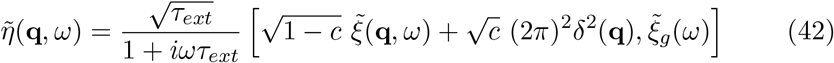

with the white noise averages,

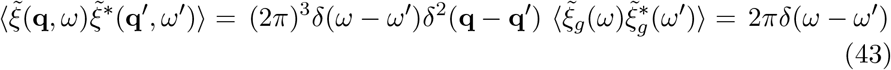

The short-hand notation *δ*^2^(**q**) has been used for the two-dimensional *δ*-function *δ*(*q*_*x*_)*δ*(*q*_*y*_). Similarly, one has for the finite size noise averages.

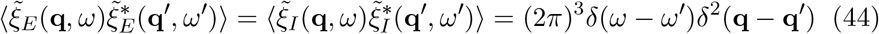

The current-current correlation function is obtained by averaging the product of the currents (Eq. (40)) over the noise with the help of Eq. (43,44). One obtains,

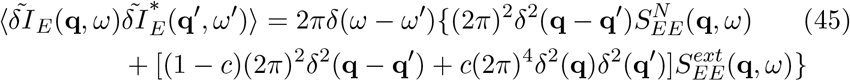

with

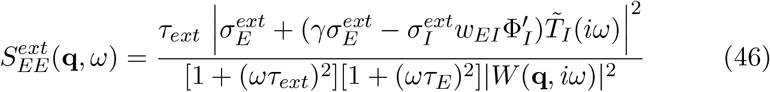

and

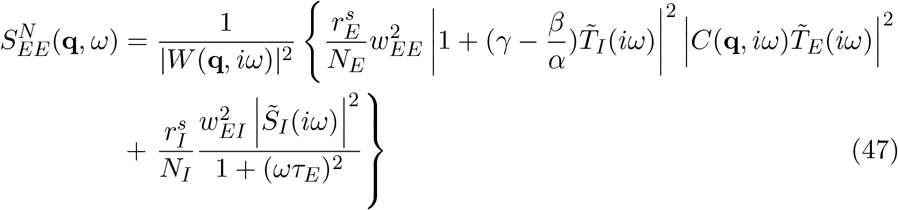

This provides the expression of the current-current correlation in real space,

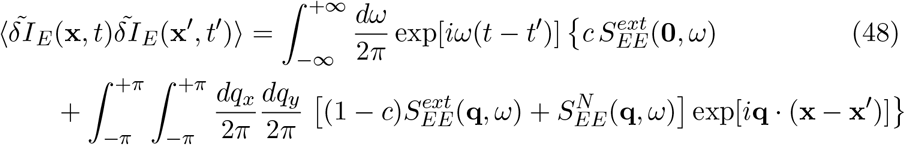

One can check that the cross-correlation is a real function, as it should, since 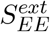 and 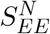 are related to their complex conjugates by 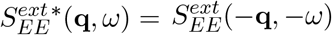 and 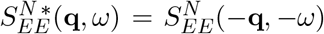, a symmetry inherited from the function *C*(**q**, *ω*) (Eq. (29)).

Taking coincident points (i.e. **x** = **x**^′^) in the current-current correlation (Eq. (48)) gives *I*_*E*_ current auto-correlation for a local module in the network. Remembering that the auto-correlation is the Fourier transform of the spectrum, this provides the spectrum *S*(*ω*) of a local module excitatory current time series,

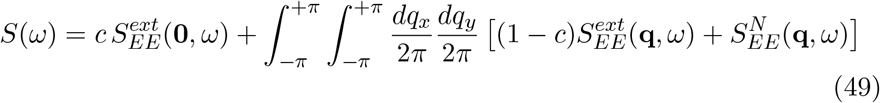

We take the local module excitatory current as a proxy for the LFP. Eq. (48) and (49) have been used to draw the theoretical lines for current-current correlations and power spectra in Fig. 2, S2, and S4.

### Simulations

The mathematical model defined in section *Model* was numerically simulated. We used a 28 × 28 grid of E-I modules. Two external layers of with E-I populations at their fixed points were added as boundary conditions. Measurements were only performed in the center square 10 × 10 grid to minimize boundary effects. A sketch of the grid is shown in Fig. S10. Model distributions and averages in all figures were obtained by performing 20 independent network simulations of 10 s simulated time each.

The reference parameters for all simulations are provided in Table 1.

Simulations were performed with a custom C program using a first-order Euler-Maruyama integration method, with a time step *dt* = 0.01 ms. Python programs were used for data analysis and to draw the figures. All simulations were performed on ECNU computer clusters.

## Acknowledgements

We are very grateful to M Denker for extensive discussions about the analysis in [12] and for allowing us to precisely compare it with our own. It is a pleasure to thank Nicolas Brunel and David Hansel for thought-provoking comments, Nicolas Brunel and Ludovica Bachschmid-Romano for sharing their unpublished work on the motor cortex and Betsy Herbert for her work and discussions during an internship on a related project. We also wish to acknowledge exchanges with Thomas Brochier and Alexa Riehle. The work of Kang Ling was supported by a CSC fellowship.

## Author contributions

Project design : JR and VH. Data analysis : LK and JR. Simulations : LK and JR. Analytical calculations : LK, JR and VH. Writing-original draft : LK and VH. Writing-review and editing : LK, JR and VH.

## Supplementary figures

**Figure S1:**
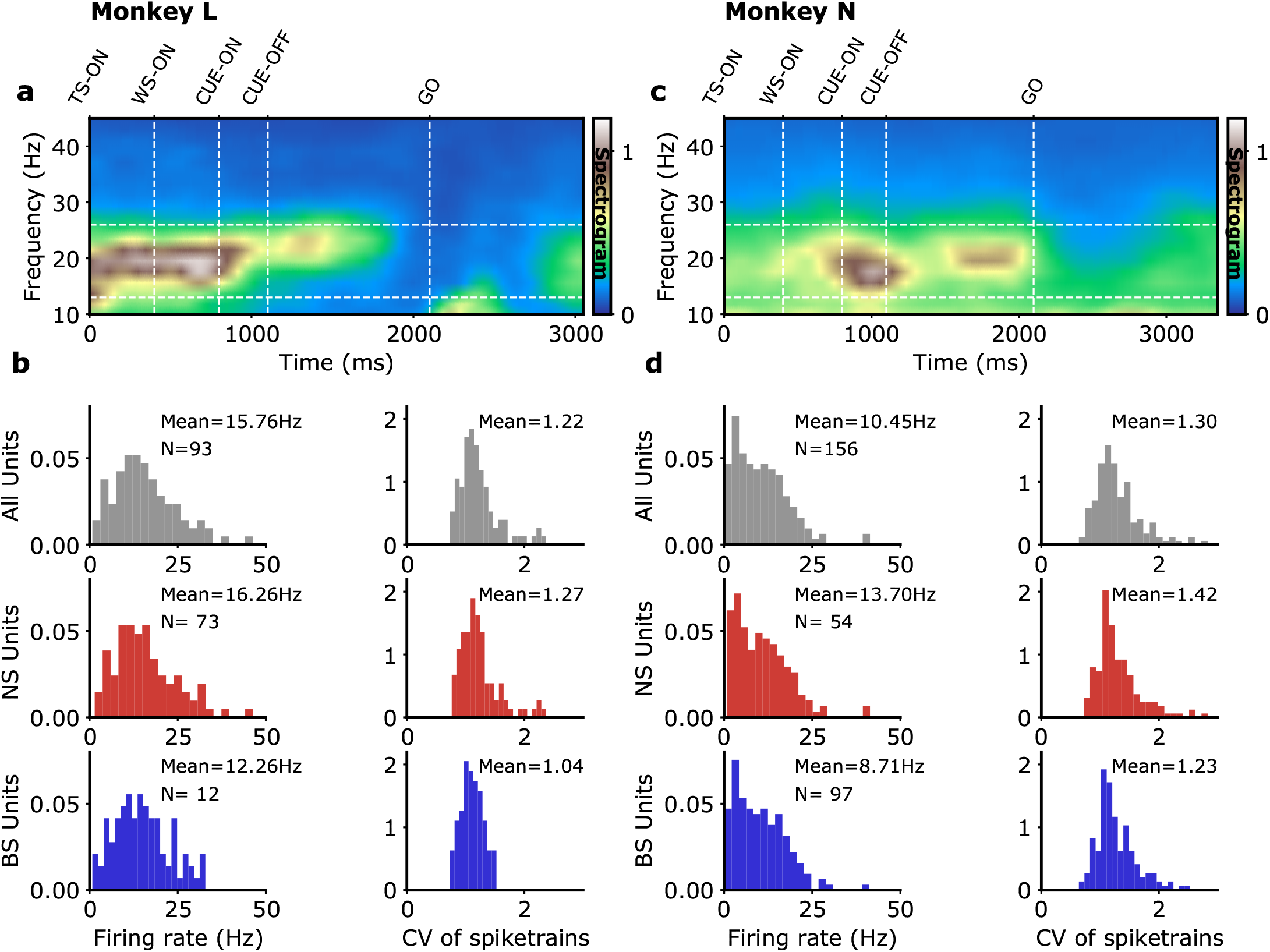
Recording data, LFP and single unit characteristics. Monkey L(a)-(b) and Monkey N (c)-(d). (a)(c) Different periods of the experiments. We focus here on the data between CUE-OFF and GO, the movement “preparatory period”. (b)(d) Characteristics of the single units spike sorted in [13] (monkey L, 93 units; monkey N, 156 units). Following Dabrowska et al.[54], spikes of width smaller than 0.4 ms were classified as Narrow Spike units (NS)/ putative interneurons (monkey L, 73 units; monkey N, 54 units) and spike of width larger than 0.41 ms were classified as Broad Spike units (BS)/ putative pyramidal cells (monkey L, 12 units; monkey N, 97 units) (units with spike width between 0.40 and 0.41 ms are not classified).

**Figure S2:**
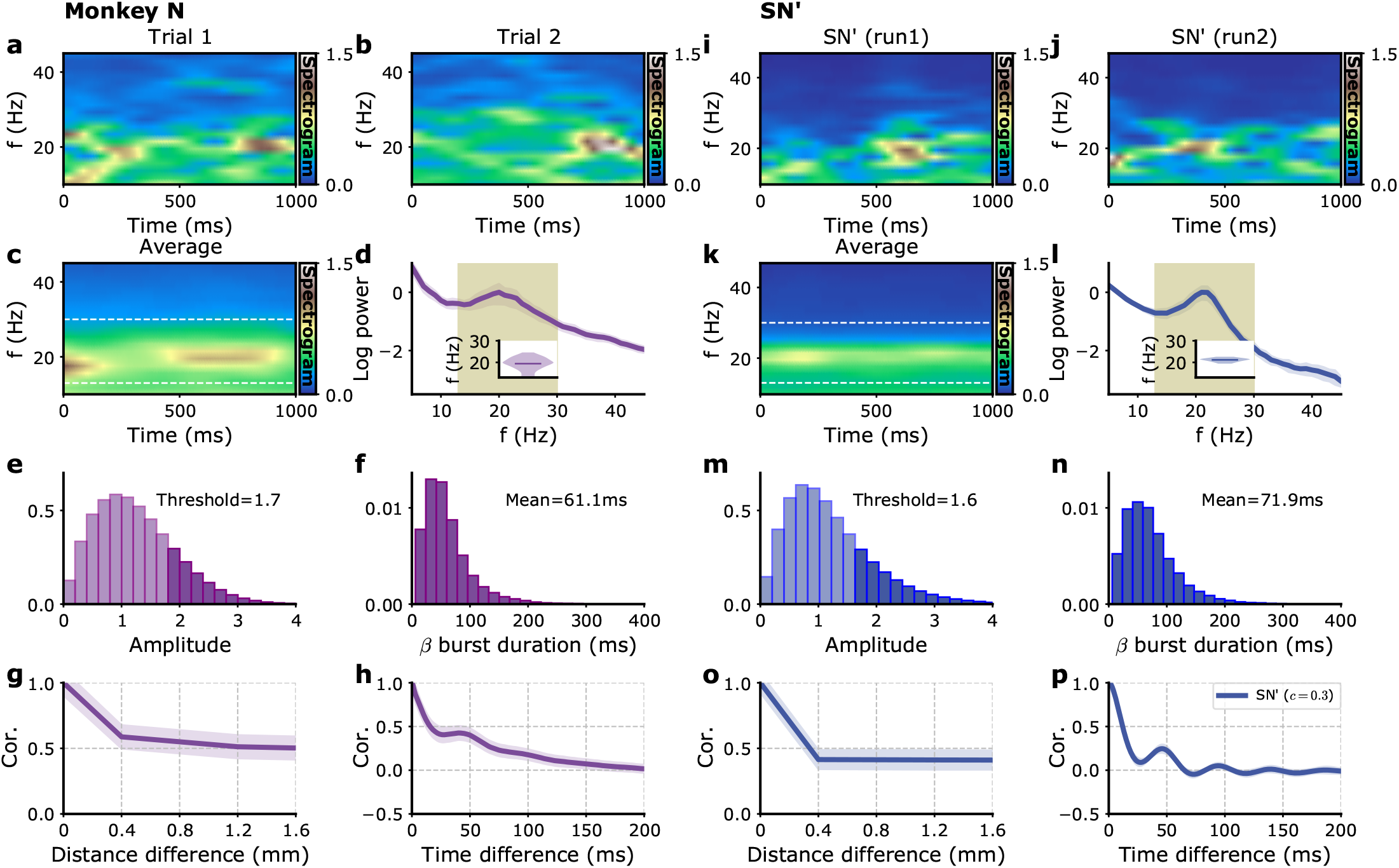
Beta bursts and power spectra for monkey N. Same as Fig. 2. Model parameters correspond to SN’ in Table 1.

**Figure S3:**
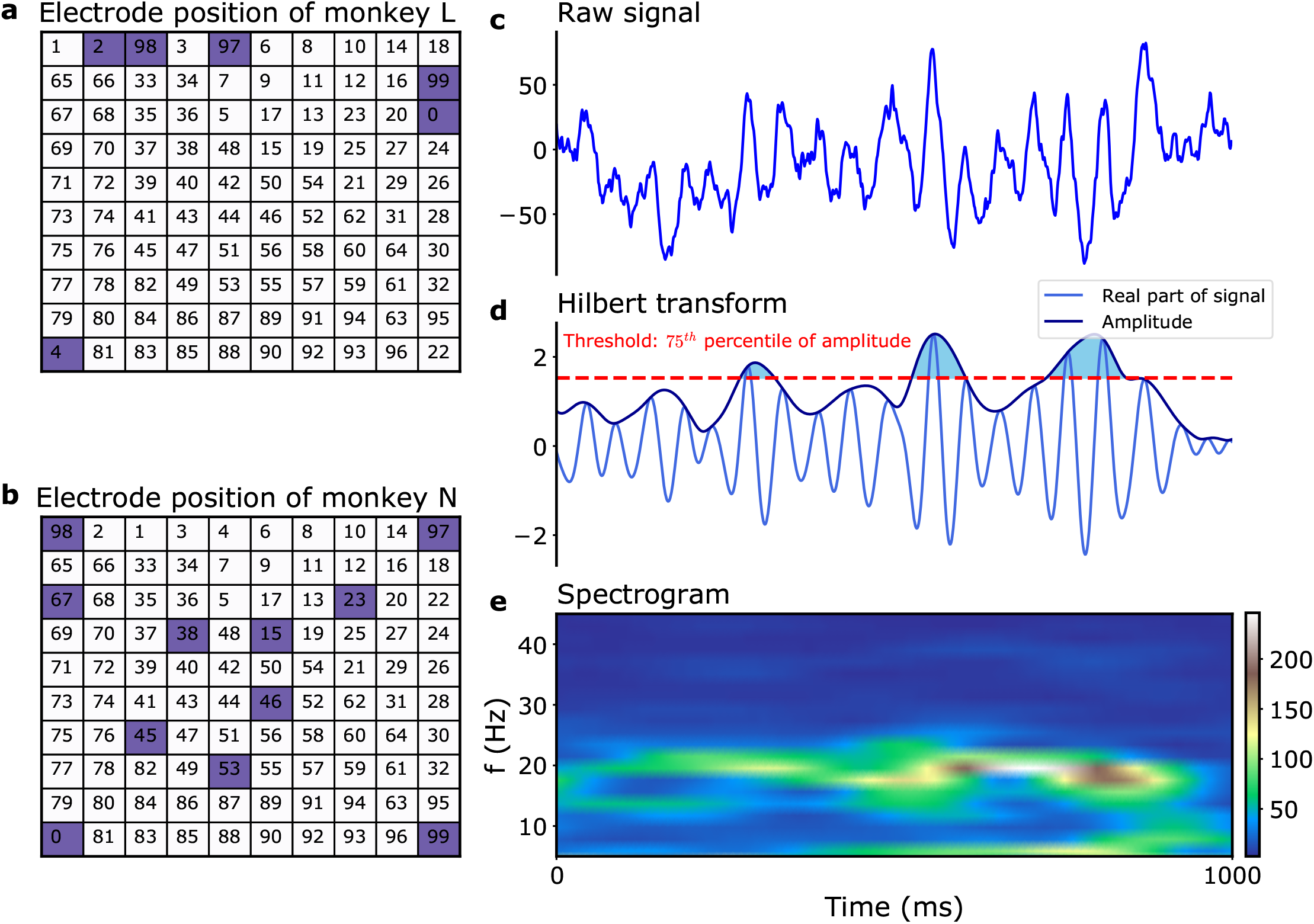
Data analysis: filtering and burst amplitude threshold. Electrode positions and numbering in [13] for (a) monkey L (b) monkey N. The blue squares indicates the positions of the dead electrodes. (c)-(e) Illustration of data analysis. (c) 1 s of single electrode signal (monkey N, electrode 7, trial 1). (d) Same signal after bandpass filtering. The amplitude threshold used to define the beta bursts is indicated (dashed red line). The corresponding beta bursts themselves correspond to the shaded blue regions (d). (e) Spectrogram of the signal in (c).

**Figure S4:**
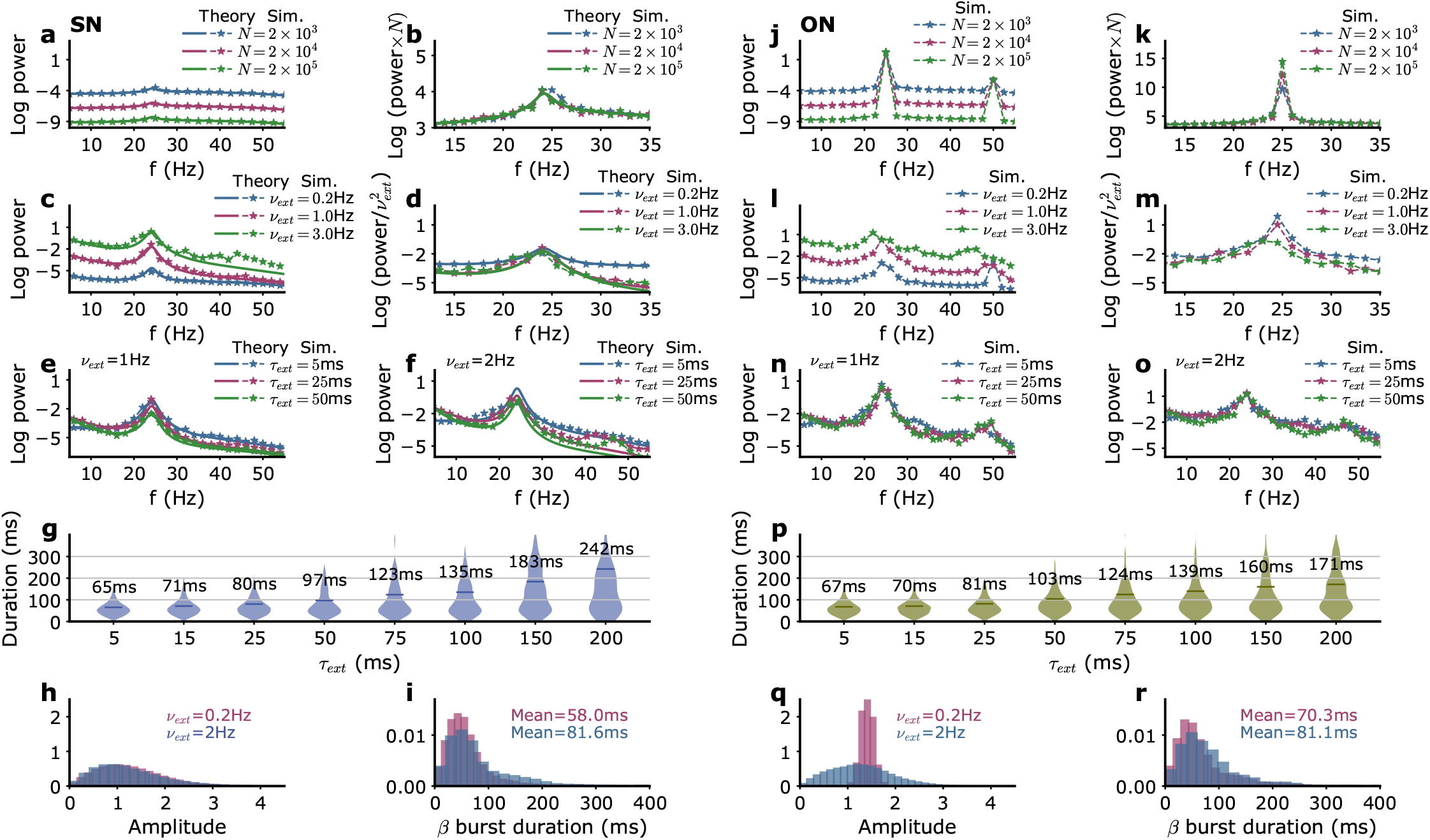
Model power spectra and beta bursts as a function of the external input parameters. (a)-(i) SN parameters, (j)-(r) ON parameters. (a)(j) Power spectra for different numbers of neurons *N* in each module corresponding to different levels of intrinsic noise without fluctuations of external inputs. For the SN parameters, the analytical formula Eq. (49) is also shown (solid lines). (b)(k) Power spectra multiplied by the number of neurons N. In this range of amplitude, the power spectra have the same shape when normalized by the amplitude of the stochastic fluctuations. (c)(l) Influence on the power spectra of the amplitude *ν*_*ext*_ of external input fluctuations. (d)(m) Power spectra divided by 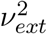. (e)(f),(n)(o) Influence on the power spectra of the correlation time *τ*_*ext*_ for two amplitudes of external input fluctuations. (g)(p) Influence on the beta burst duration of the correlation time *τ*_*ext*_ of external input fluctuations. Distribution of beta burst amplitudes (h)(q) and durations (i)(r) for two amplitudes *ν*_*ext*_ of external input fluctuations. Parameters of the input that are not explicitly varied as well as the network parameters for models SN and ON are given in Table 1.

**Figure S5:**
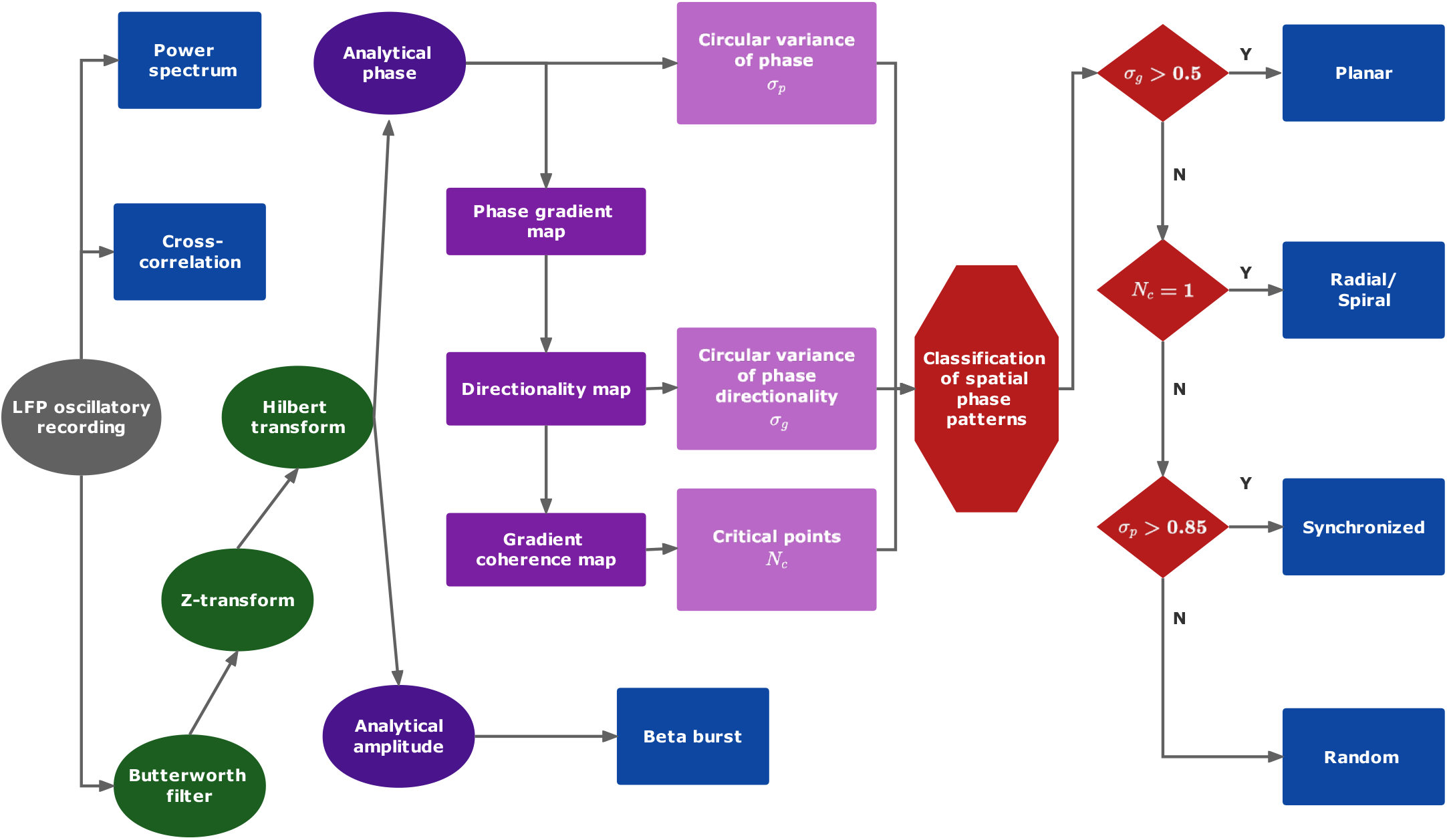
Data analysis protocol. See *Methods* for a description of the different steps.

**Figure S6:**
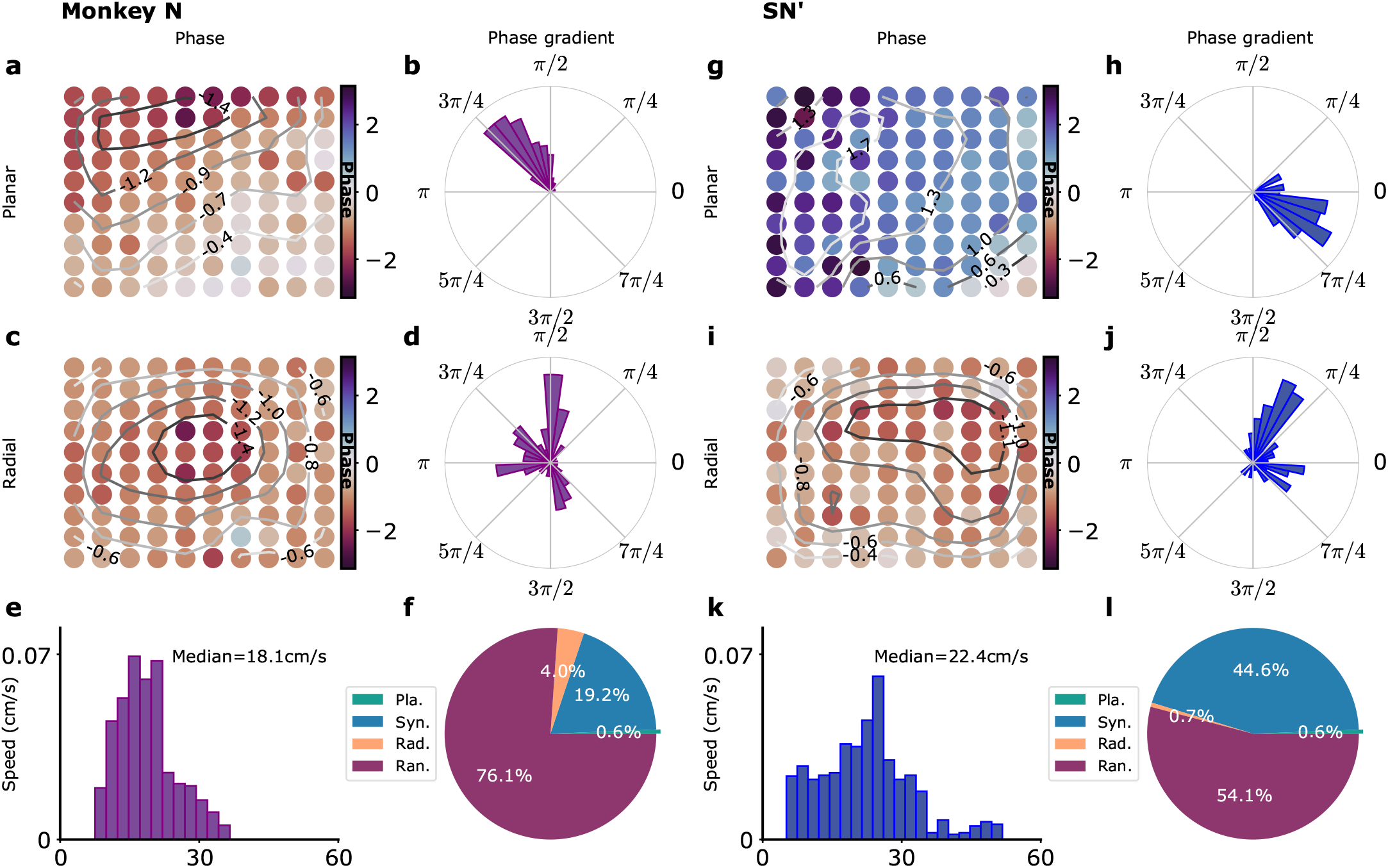
Waves in model simulations and in recordings for monkey N. Same as Fig. 4. (b) *σ*_*g*_ = 0.59. (d) *σ*_*g*_ = 0.10. (h) *σ*_*g*_ = 0.53. (j) *σ*_*g*_ = 0.23.

**Figure S7:**
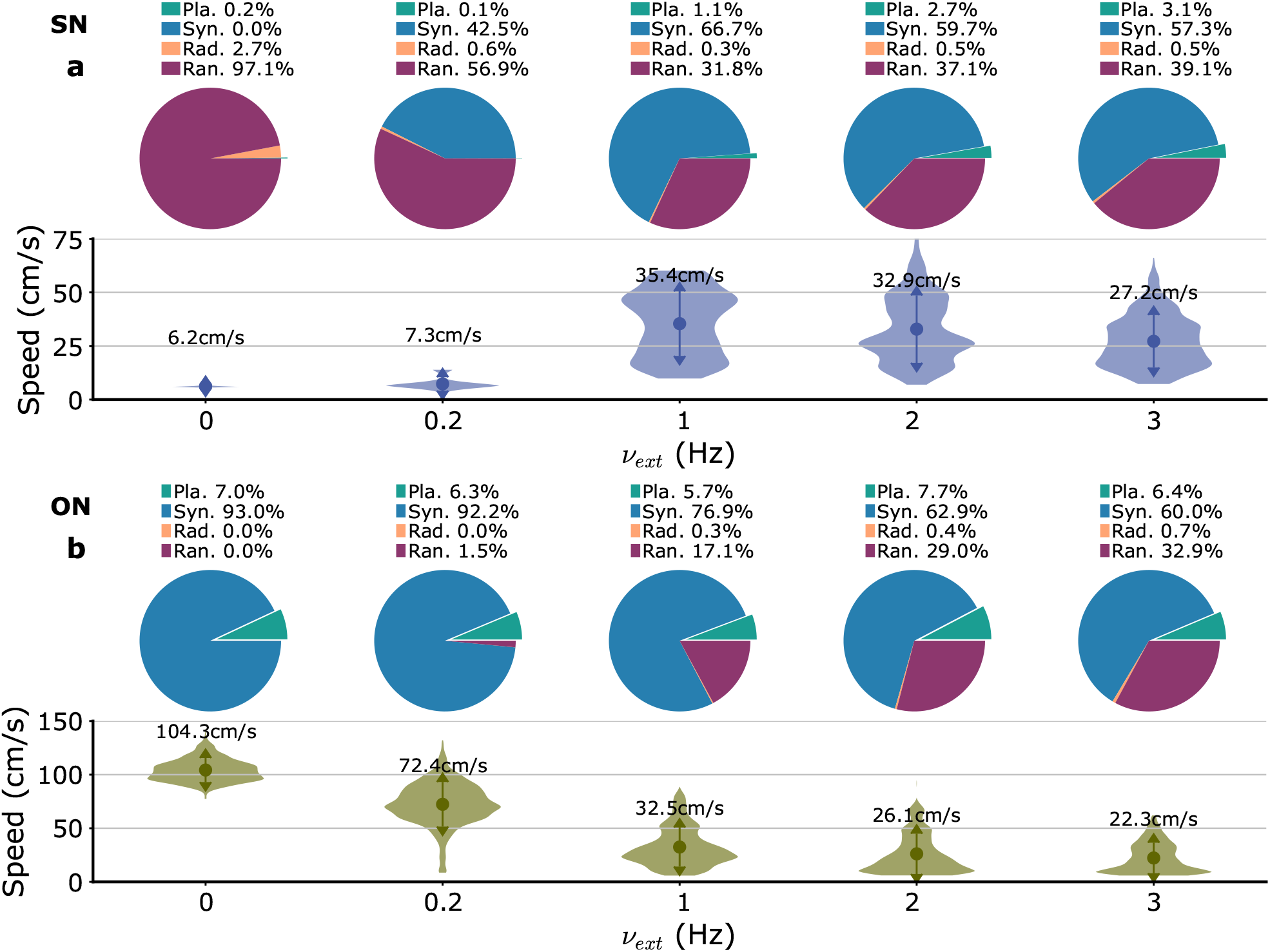
Variation of wave type distribution and planar wave speed with the amplitude of the external input fluctuations (*ν*_*ext*_). (a) SN parameters. (b) ON parameters.

**Figure S8:**
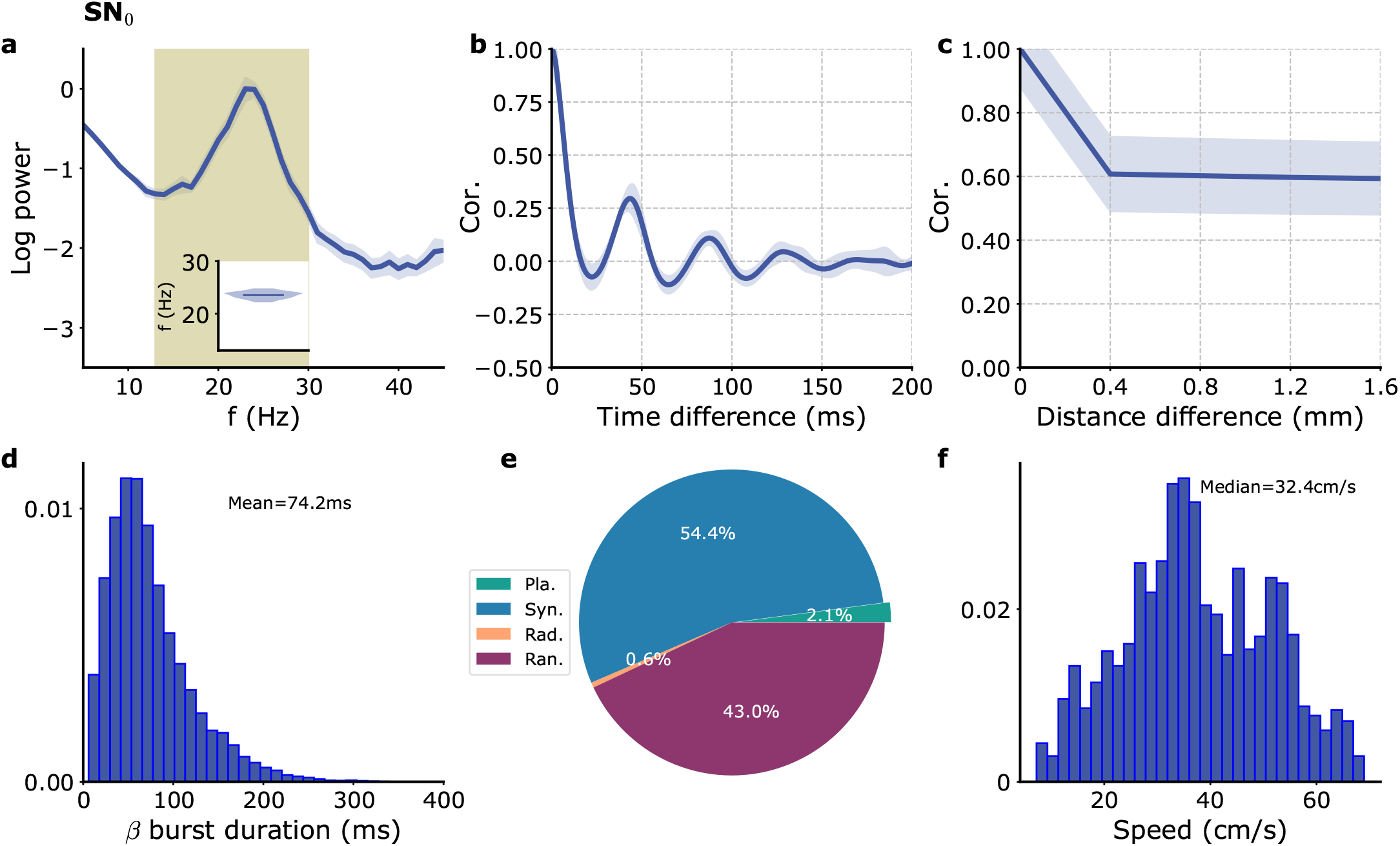
Model with no propagation delay. (a) Single module *I*_*E*_ power spectrum. (b) Single module *I*_*E*_ auto correlation. (c) *I*_*E*_ cross-correlation as a function of module distance. (d) Distribution of beta burst duration. (e) Proportion of different wave types. (f) Distribution of speeds of planar waves. Model parameters correspond to SN_0_ in Table 1.

**Figure S9:**
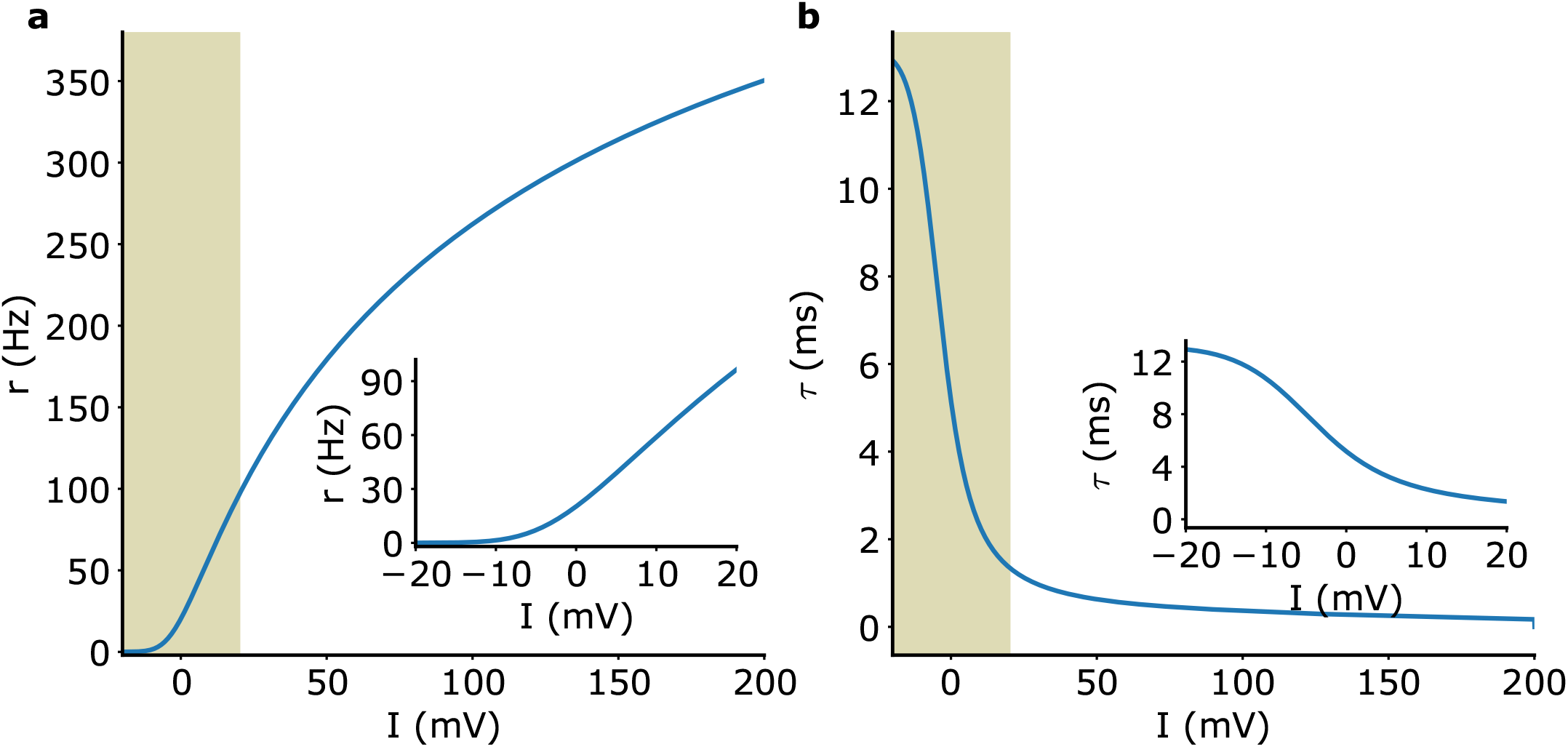
Rate model f-I curve and adaptive time scale. (a) f-I curve. Insert : zoom on the 0 − 100 Hz part. (b) Adaptive time scale. The data corresponding to these curves are provided in *Source data*.

**Figure S10:**
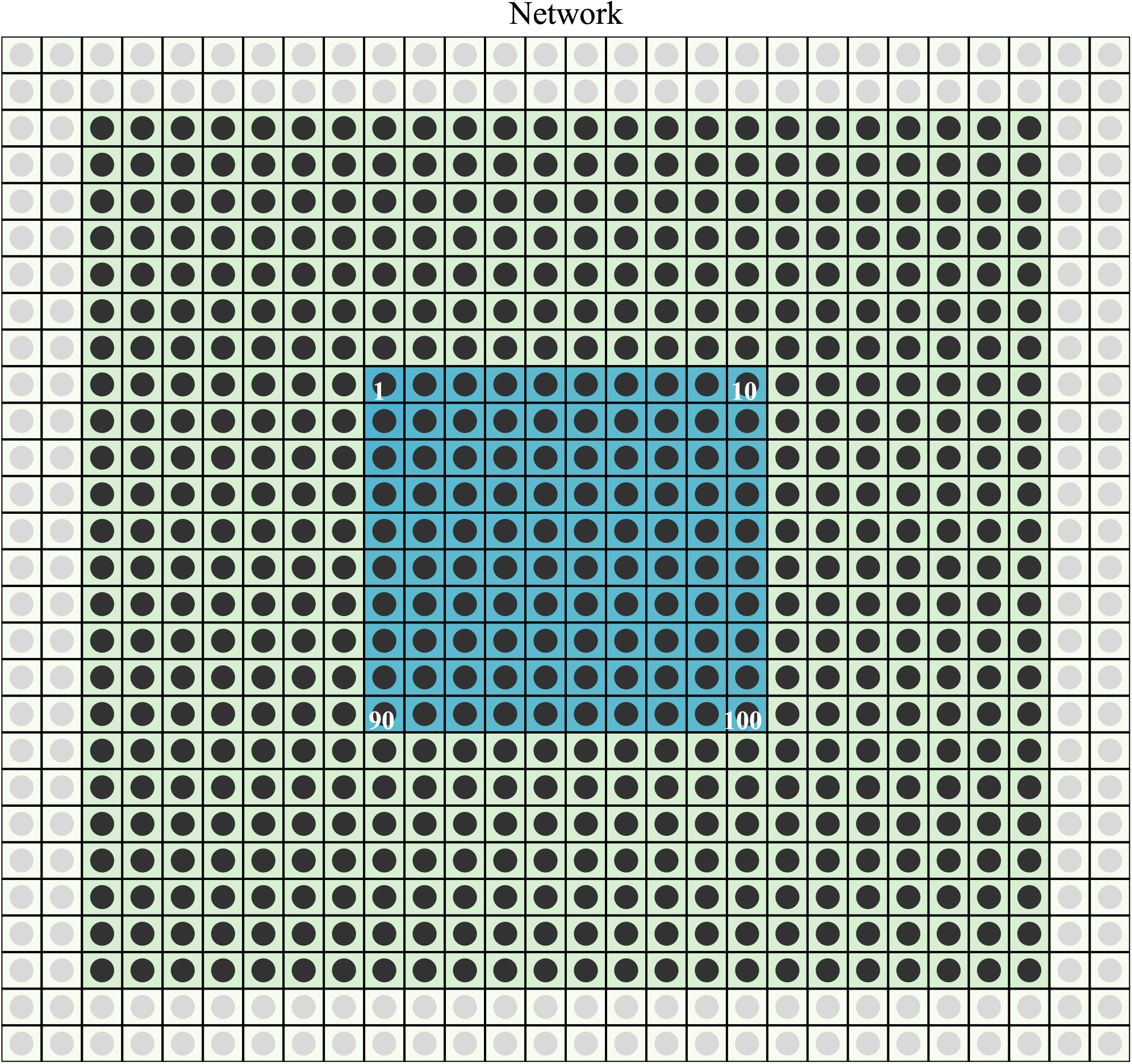
Numerical simulation grid. Different E-I modules (disk) are placed at the center of the different squares. The discharge rates of the E-I populations in two most external layers (gray disks) are fixed at their steady state values. The other modules (black disks) in the 24 × 24 central array are simulated. Only the modules in the 10 × 10 central array (blue squares) are used for the different signal measurements.

